# A Spinal Circuit for Modular Gating of Organ Somatosensation

**DOI:** 10.1101/2025.08.27.672386

**Authors:** Yufen Zhang, Fang Liu, Lite Yang, Hannah J Hahm, Monique R Heitmeier, Takao Okuda, Masahiro Kawatani, Jahnia T Harrigan, Erica C Morrison-Rodriguez, Anna Petukhova, Zhaotong Lu, Harry A Heitmeier, Vijay K Samineni

**Affiliations:** Department of Anesthesiology, Washington University Pain Center, Washington University School of Medicine, St. Louis, MO, United States; Neuroscience Graduate Program, Division of Biology & Biomedical Sciences, Washington University School of Medicine, St. Louis, MO, United States

**Author notes:** Co-first authors (Yufen Zhang, Fang Liu).

## Abstract

End-organs such as the bladder rely on a delicate balance between internal urgency and voluntary restraint. However, the specific spinal circuits coordinating these functions remain poorly defined. Here, we identify a genetically defined population of lumbosacral spinal interneurons marked by *Trpc4* expression that gate bladder sensory input and regulate micturition reflexes. Single-cell transcriptomics and in situ physiology reveal molecularly distinct *Trpc4*⁺ subtypes. Functional manipulation reveals that *Trpc4*⁺ neurons are essential for coordinating bladder-sphincter activity and gating visceral pain. Their ablation leads to bladder hypersensitivity and voiding dysfunction, while targeted activation reverses these maladaptive states. Circuit tracing reveals convergence of primary afferent and descending brainstem inputs onto *Trpc4*⁺ neurons. These findings establish *Trpc4*⁺ interneurons are critical for bladder sensory-motor integration and extend classical spinal gating models to encompass visceral pain and organ reflex control.

## Main Text

Among visceral organs, the bladder is uniquely subject to voluntary control, requiring coordinated activity between the peripheral and central nervous systems ^1–3^. While peripheral mechanisms of bladder sensation have been increasingly well characterized ^4–9^, the central circuit logic and molecular identity of spinal neurons that integrate bladder input and orchestrate voiding reflex remain largely undefined. The lumbosacral spinal cord (SC), particularly its medial region, the dorsal gray commissure (DGC), is a critical node for processing sensory input from the lower urinary tract receiving convergent input from both innocuous and noxious bladder afferents, highlighting its dual role in bladder sensation and pain ^10–15^. Electrophysiological studies found that the DGC neurons encode non-nociceptive responses related to bladder filling, voiding, and noxious distension-induced pain ^16–18^, suggesting that the lumbosacral cord is an integral part of neural mechanisms that govern bladder somatosensation ^18–20^. Despite this foundational knowledge, the specific cell types and circuit logic that govern bladder reflexes, nociceptive processing, and voluntary control remain poorly defined. Bridging this gap is essential for understanding how spinal circuits translate sensory input into context-dependent motor outputs ranging from routine voiding to pathological states such as bladder pain syndromes and voiding dysfunction. Here, we begin to address this gap by molecularly, physiologically, and anatomically defining the spinal neurons that regulate bladder sensation.

## Molecular Profiling Reveals *Trpc4***⁺** Neuron Enrichment in Lumbosacral SC

To gain genetic access to molecularly distinct lumbosacral spinal populations, we performed single-nucleus RNA sequencing (snRNA-seq) on micro dissected DGC (n = 3) and whole lumbosacral SC (n = 4) samples. This enabled the construction of a lumbosacral-focused single-cell atlas and comparison with existing atlases from more rostral spinal levels ^21^. After quality control, 51,352 nuclei (27,295 neuronal; 24,057 non-neuronal) were retained, with minimal batch effects and overlapping UMAP distributions across replicates (Extended Fig. 1a, b). Clustering identified 32 neuronal subtypes (Fig. 1a, Extended Data Fig.1e) and 7 non-neuronal classes (Extended Data Fig.1c, d).

**Fig. 1.**
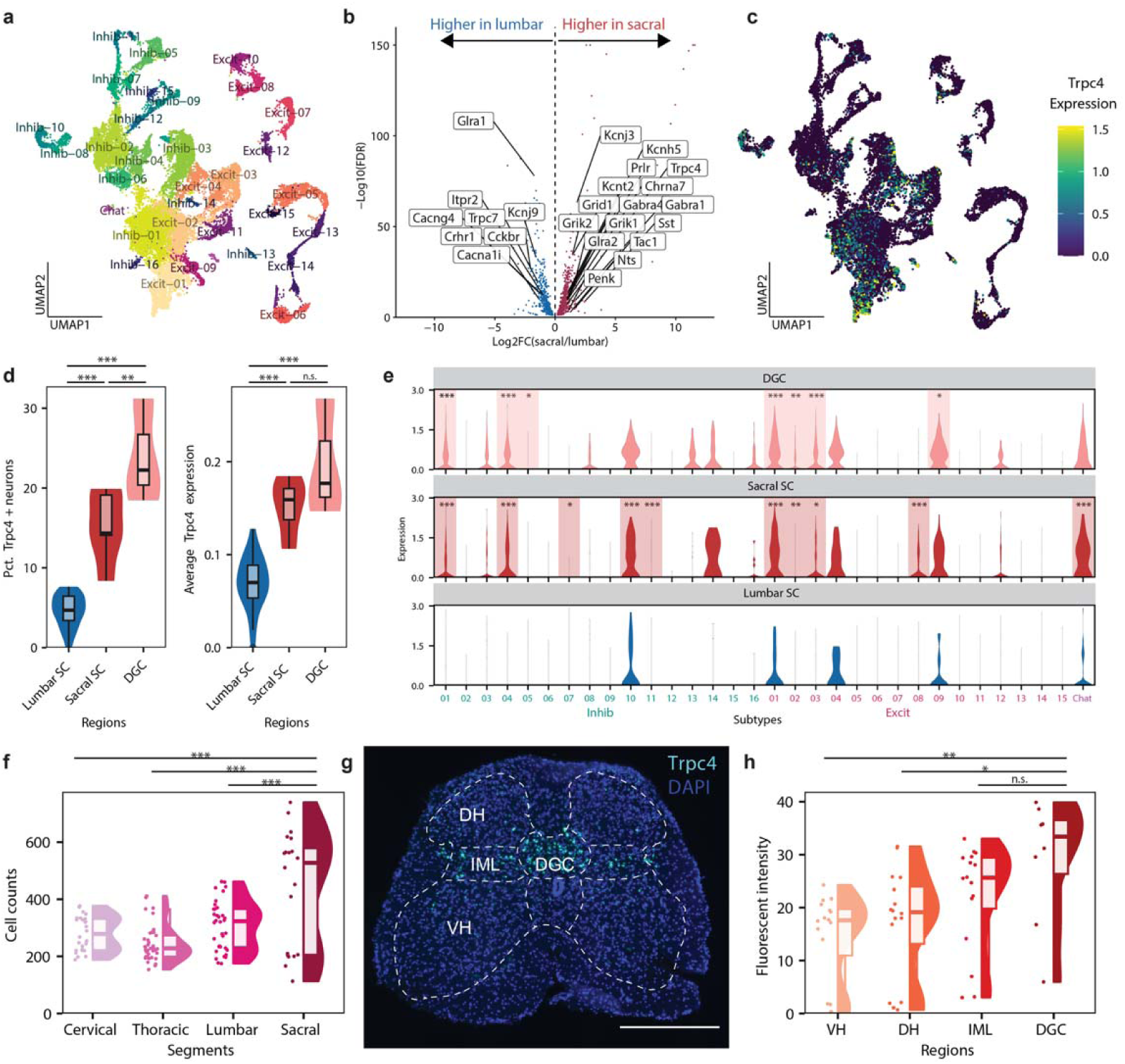
Molecular profiling reveals *Trpc4⁺* neuron enrichment in lumbosacral SC. (a) snRNA-seq UMAP showing the neuronal nuclei from sacral SC and DGC. Nuclei are colored by neuronal subtypes. (b) Volcano plot showing the differentially expressed genes between lumbar SC neurons from previously reported snRNA-seq data and sacral SC neurons reported in this study. Labels denote significantly differentially expressed genes (Log2FC>1, FDR<0.05) of neuropeptides, peptide receptors, G protein-coupled receptors, and ion channels. (c) UMAP showing the expression of *Trpc4* in neuronal nuclei from sacral SC and DGC. (d) Bar plots and violin plots showing the *Trpc4* expression in neurons from lumbar SC, sacral SC, and DGC in snRNA-seq data. **p<0.01, ***p<0.001, Post-hoc Tukey’s HSD following ANOVA. (e) Violin plots showing the expression of *Trpc4* in each neuronal subtype from lumbar SC, sacral SC, and DGC snRNA-seq data. *p<0.05, **p<0.01, ***p<0.001, Post-hoc Tukey’s HSD following ANOVA for each subtype. (f) Cell counts from spinal segments showing *Trpc4* expression. ***p<0.001, Post-hoc Tukey’s HSD following ANOVA for each subtype. (g) Representative fISH image showing *Trpc4* expression in the lumbosacral SC. (h). Cell counts lumbosacral SC showing *Trpc4* expression distribution in subdivisions of SC. *p<0.05, **p<0.01, Post-hoc Tukey’s HSD following ANOVA for each subtype.

Most neuronal subtypes aligned with known spinal classes, supporting overarching segmental continuity in cell type organization (Extended Fig. 2a-d). However, 1,132 genes were differentially enriched in lumbosacral neurons, including 16 neuropeptides, receptors, and ion channels (Fig.1b). Among these, Trpc4 was selectively enriched in the lumbosacral SC and especially the DGC (Fig. 1c–d). *Trpc4* expression was elevated in 10 lumbosacral neuronal types and 7 DGC clusters relative to their lumbar counterparts (Fig. 1e), a pattern corroborated by the Allen Brain Atlas (Extended Fig. 2e) and Fluorescent in situ hybridization (fISH) for *Trpc4* in SC (Fig.1f, Extended Data Fig. Extended Data Fig.2f). fISH analysis confirmed spatially restricted *Trpc4* expression in the DGC, dorsal horn (DH), and intermediolateral column (IML) of lumbosacral SC (Fig.1g, h). Given the DGC’s known role in receiving visceral afferents, which in turn participate in both nociceptive signaling and reflex circuits governing micturition and pelvic floor control ^12,14,22,23^, *Trpc4*⁺ neurons may provide genetic access to lumbosacral sensorimotor circuits governing bladder pain and voiding reflex.

## DGC*^Trpc4^* Neuron Ablation Impairs Bladder Sensory Gating

To assess whether DGC*^Trpc4^* lumbosacral neurons contribute to bladder sensory integration and reflex control, we genetically ablated these neurons. AAV1-DIO-taCasp3 was injected into the lumbosacral SC of *Trpc4*-Cre::Ai14 mice to express Cre-dependent caspase-3, while controls received AAV5-DIO-eYFP. Injections were targeted to the DGC, as confirmed histologically (Fig. 2a, b). taCasp3 resulted in ∼90% reduction of *Trpc4*⁺ neurons relative to controls (Fig. 2c), enabling functional dissection of their roles. To evaluate the contribution of DGC*^Trpc4^*neurons to bladder nociception, we recorded visceromotor reflexes (VMRs) during graded bladder distension (20–80 mmHg) (Fig. 2d). Control mice showed a pressure-dependent VMR increase, *Trpc4*-ablated mice exhibited exaggerated responses at all intensities (Fig. 2e-f), suggesting DGC*^Trpc4^* neurons gate bladder sensory input analogous to prior work that showed lumbosacral SC neurons that are inhibited by the noxious colorectal distensions ^24^. Ablated mice also showed enhanced mechanical sensitivity to abdominal von Frey stimulation, consistent with referred visceral hypersensitivity (Fig. 2g). However, withdrawal thresholds to mechanical and thermal stimuli applied to the hindpaw remained unchanged (Extended Fig. 3a-d), indicating that *Trpc4*⁺ neurons modulate pelvic-specific sensory input in a segment- and modality-selective manner.

**Fig. 2.**
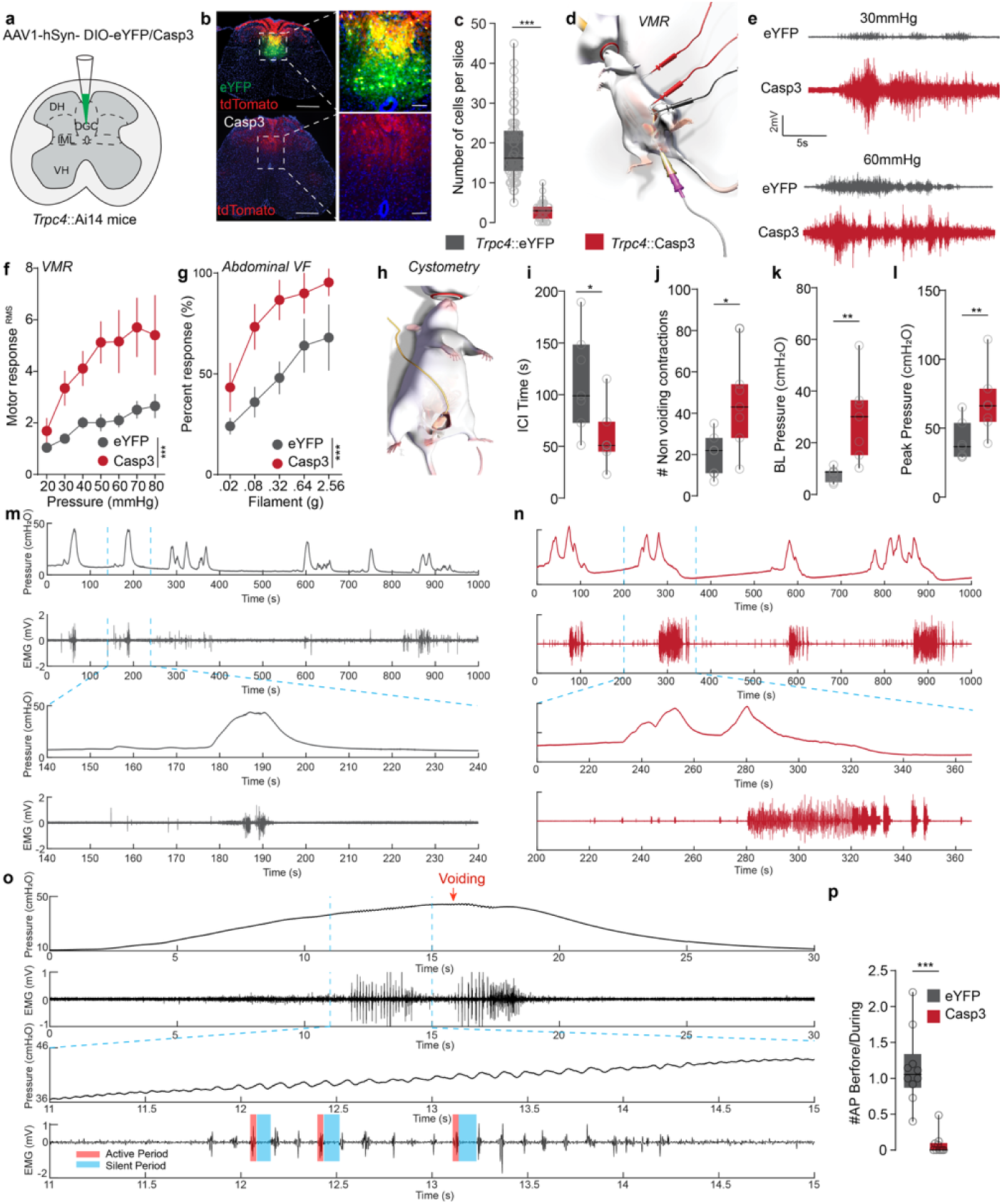
DGC*^Trpc4^* neurons are necessary for bladder sensory gating and sphincter coordination. (a) Schematic of AAV1-hSyn-DIO-eYFP or -Casp3 injection targeting the DGC of *Trpc4*::Ai14 mice. (b) Representative SC sections showing *Trpc4⁺* neurons after virus injection in *Trpc4*::Ai14 mice. Scale bars, 500 µm (overview), 100 µm (zoom). (c) Quantification of *Trpc4⁺* neurons remaining after eYFP or Casp3 injection (***P < 0.001, Mann–Whitney U test). (d) Schematic of VMR to bladder distension in anesthetized mice. (e) Representative EMG traces from eYFP and Casp3 mice during bladder distension at 30 and 60 mmHg. (f) Graded VMR responses to bladder distension pressure show increased responses in Casp3 mice compared to eYFP controls (***P < 0.001, two-way ANOVA). (g) Abdominal mechanical hypersensitivity assessed by von Frey stimulation reveals increased withdrawal responses in Casp3 mice (*P < 0.05, two-way ANOVA). (h) Illustration of cystometry setup in anesthetized mice. (i–l) Quantification of bladder function shows shortened ICIs (i), more NVCs (j), elevated baseline pressure (k), and elevated peak pressure (l) in Casp3 mice (*P < 0.05, *P < 0.05, **P < 0.01, **P < 0.01, unpaired t test). (m and n) Representative EUS-CMG traces show simultaneous EUS-EMG activity during micturition in eYFP and Casp3 mice. (o) Expanded EUS-EMG traces illustrate bursting activity before and during voiding, with red and blue boxes denoting active and silent periods, respectively. (p) Quantification of EUS active periods during pre-void and void phases shows decreased EUS activity in pre-void phase in Casp3 mice (***P < 0.001, Mann–Whitney U test).

## Loss of DGC*^Trpc4^* Neurons Disrupts Bladder–Sphincter Coordination

Electrical stimulation of the DGC is shown to regulate the voiding reflex ^25^, yet the specific cell types mediating this function remain unidentified. We next asked whether DGC*^Trpc4^* neurons contribute to micturition control (Fig. 2h). Urodynamic measurements revealed that *Trpc4*-ablated mice displayed shortened intercontraction intervals (ICIs), more frequent non-voiding contractions (NVCs), and elevated bladder pressures (Fig. 2i–k, Extended Data Fig.3e, f) resembling cystitis-associated bladder dysfunction ^26,27^. To directly assess the effects of DGC*^Trpc4^* neuron ablation on urinary muscle coordination, we performed simultaneous external urethral sphincter (EUS) electromyography (EMG) alongside cystometric recordings. In control mice, increases in bladder pressure during voiding were tightly coupled with high-frequency EUS bursting, followed by bladder pressure drops during voiding (Fig. 2m). These EUS bursts displayed organized transitions between high-activity active periods (APs) and silent periods (SPs), consistent with normal detrusor-sphincter synergy (Fig. 2o). In contrast, *Trpc4*-ablated mice displayed atypical EUS bursting during non-voiding bladder contraction bursts that failed to trigger a corresponding reduction in bladder pressure (Fig. 2n). Quantitative analysis revealed a significant reduction in EUS APs surrounding bladder contractions in *Trpc4*-ablated animals compared to controls (Fig. 2p), indicating impaired EUS recruitment.

Given prior evidence that DGC GABAergic interneurons regulate EUS activity via inhibitory input to motoneurons in Onuf’s nucleus ^28–31^, we asked whether DGC*^Trpc4^* neurons directly innervate motoneurons. We co-injected CTB into the EUS to retrogradely label sphincter-projecting motoneurons and expressed tdTomato-T2A-Syp-eGFP in DGC*^Trpc4^*neurons. This allowed visualization of synaptic terminals versus fibers, revealing dense bilateral projections to the DH, IML, and ventral horn key regions for nociceptive processing and urinary reflex control (Extended Data Fig.3g-h). We observed Syp-eGFP⁺ puncta from DGC*^Trpc4^* neurons in close apposition to CTB-labeled motoneurons in Onuf’s nucleus (Extended Data Fig.3i), providing anatomical evidence for direct connectivity. Notably, DGC*^Trpc4+^* axons were confined to the spinal cord, with no evidence of supraspinal projections (Extended Data Fig.3j-n), confirming their identity as spinal interneurons. Together, these results demonstrate that DGC*^Trpc4^* neurons are essential for synchronizing detrusor and EUS activity during voiding. Their loss disrupts the coordination required for effective voiding and promotes hyperreflexic bladder activity. The direct anatomical connection to sphincter-controlling motoneurons further supports a mechanistic model in which DGC*^Trpc4^* neurons act as key integrators in spinal microcircuits that govern bladder-sphincter coordination, enabling proper execution of the micturition reflex.

## Classification *DGC^Trpc4^* Neuronal Physiological Diversity

We examined if pathological conditions such as cystitis, which induce bladder inflammation, pain, and dysregulated voiding, alter the activity of DGC*^Trpc4^* interneurons. To this end, first, we catalogued the physiological diversity in the DGC*^Trpc4^* interneurons, as prior work demonstrated heterogeneity of DGC neurons^32^. We performed targeted whole-cell patch-clamp recordings and biocytin fills in acute lumbosacral SC slices from *Trpc4*-Cre;Ai14/Ai213 mice (Fig. 1d, Extended Data Fig.2a), recording from 122 GFP⁺/tdTomato⁺ neurons. Analysis revealed five firing phenotypes: single-spiking, delayed, gap, phasic, and tonic (Fig. 3a–c, Extended Data Fig.4a-c), resembling previously described DGC neurons; others matched profiles of known spinal interneuron classes ^32–34^. Hierarchical clustering based on intrinsic properties confirmed these as distinct electrophysiological subtypes (E-types), validating our manual classification (Fig. 3d). Morphological reconstruction of biocytin-filled neurons showed distinct somatic shapes, dendritic branching, and axonal projections were evident across E-types (Fig. 3e, Extended Data Fig.4e). Single-spiking and phasic neurons showed axons restricted to the DGC, while delayed, gap, and tonic-firing types extended collaterals into the IML. Whereas gap-firing neurons projected to the DH and tonic neurons had the most elaborate axonal trajectories to ventral SC, suggesting broader circuit engagement. We integrated morphometrics with firing data for multimodal clustering, which revealed two spatially distinct DGC*^Trpc4^* clusters: one comprising

**Fig. 3.**
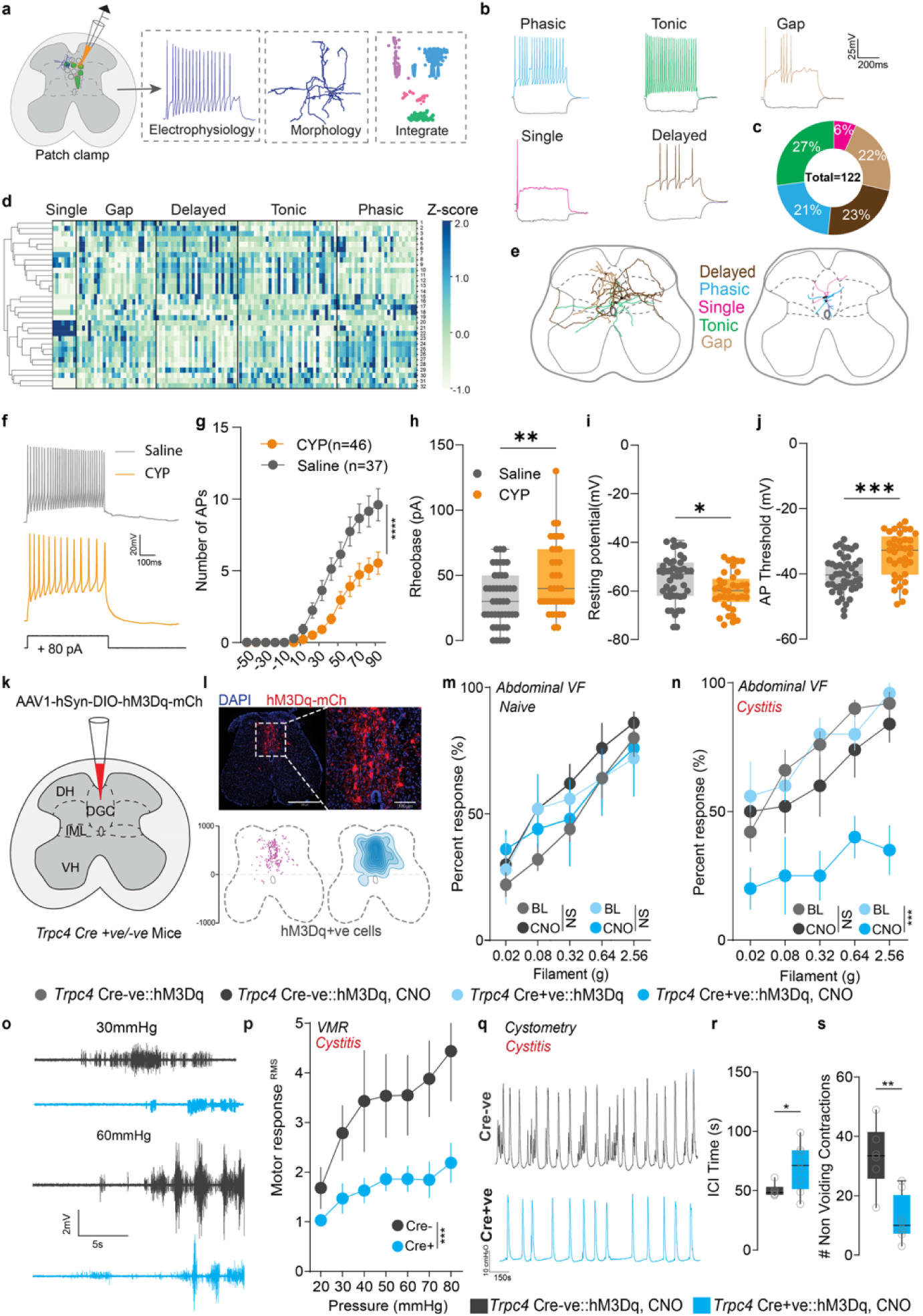
Dampened DGC*^Trpc4^* neuronal activity causes bladder pain and voiding dysfunction. (a) Experimental workflow of patch-clamp recordings and morphological reconstruction of *Trpc4*⁺ DGC neurons in the lumbosacral SC. (b) Representative firing patterns identified and their distribution (c) in *Trpc4*⁺ DGC neurons. (d) Heatmap showing hierarchical clustering of electrophysiological parameters (Extended Data Table1) recorded from *Trpc4*⁺ DGC neurons. (e) Schematic showing anatomical locations of morphologically reconstructed neurons across lumbosacral SC identified by E-types. (f) Representative traces from *Trpc4*⁺ neurons of saline- and CYP-treated mice. (g–j) Cystitis significantly reduced *Trpc4*⁺ neuron excitability, deceased number of action potentials (g, ****P < 0.0001, two-way ANOVA), increased rheobase (h, **P < 0.01, unpaired t test), hyperpolarized resting membrane potential (i, *P < 0.05, unpaired t test), and elevated AP threshold (j, ***P < 0.001, unpaired t test). (k) Schematic of AAV1-hSyn-DIO-hM3Dq-mCherry injection into the DGC of *Trpc4*::Cre⁺ and Cre⁻ mice. (l) Representative spinal cord section showing localization and spatial mapping of hM3Dq⁺ neurons in the DGC. (m) Chemogenetic activation of *Trpc4+* DGC neurons had no effect on abdominal von Frey testing. (P = 0.6056, two-way ANOVA). (n) Chemogenetic activation of *Trpc4+* DGC neurons significantly reduced mechanical hypersensitivity in cystitis mice compared to baseline and Cre-ve controls. (n, ***P < 0.001, two-way ANOVA). (o) Representative EMG recordings from Cre⁺ and Cre⁻ mice to bladder distension (30 and 60 mmHg). (p) VMR to graded bladder distension pressures is attenuated in CNO-treated Cre⁺ cystitis mice (***P < 0.001, two-way ANOVA) compared to Cre-ve. (q) Cystometry traces show improved voiding function following chemogenetic activation of *Trpc4*⁺ neurons. (r) ICI is prolonged, (s) NVCs are reduced in Cre⁺ mice after CNO treatment (*P < 0.05, **P < 0.01, unpaired t test). NS, not significant.

DGC-restricted phasic/single-spiking neurons, and another of IML-projecting delayed, gap, and tonic types (Extended Data Fig.4d). These results suggest DGC*^Trpc4^*interneurons form modular microcircuits, with subtype-specific spinal projection domains associated with distinct sensory-motor functions. Finally, prior efforts in the SC ^35^ have shown that DH E-types can be classified on the expression of excitatory and inhibitory neurotransmitters. We performed patch-seq on a subset of electrophysiologically characterized neurons from *Trpc4*-Cre;Ai14/Ai213 mice to identify neurotransmitter types (Extended Data Fig.5a-c). Consistent with previous observations in DH circuitry, tonic-firing neurons exhibited enriched expression of inhibitory marker (*Gad2)* aligned with inhibitory molecular signatures (Extended Data Fig.5d). In contrast, delayed- and gap-firing neurons showed transcriptomic features indicative of excitatory cell types, including expression of *Slc17a6* (Extended Data Fig.5d), while phasic-and single-firing neurons composed of a heterogeneous population of excitatory and inhibitory markers. To independently validate these molecular identities, we performed fISH for *Slc32a1* (*Vgat*, inhibitory) and *Slc17a6* (*Vglut2*, excitatory) on SC sections. Quantitative analysis revealed that approximately 36% of *Slc32a1*⁺ and over 48% of *Slc17a6*⁺ neurons co-expressed *Trpc4* (Extended Data Fig.5e), confirming that DGC*^Trpc4^ inter*neurons represent a heterogeneous population composed of both inhibitory and excitatory subtypes. This molecular dichotomy was further supported by the differential expression of ligand- and voltage-gated ion channels across subtypes (Extended Data Fig.5f). Together, these data demonstrate that DGC*^Trpc4^* interneurons in the lumbosacral SC are transcriptionally, physiologically, and morphologically diverse.

Finally, we assessed whether cystitis modulates the intrinsic excitability and firing properties of DGC*^Trpc4^* interneurons under pathological conditions. Whole-cell recordings from DGC*^Trpc4^ inter*neurons revealed that CYP-induced cystitis significantly alters their intrinsic excitability. Compared to saline controls, DGC*^Trpc4^ inter*neurons from CYP-treated mice exhibited decreased firing in response to depolarizing current steps (Fig. 3f-g, Extended Data Fig.6h-l), along with a significantly higher rheobase, more hyperpolarized resting membrane potential, and elevated action potential threshold (Fig. 3h-j. Extended Data Fig.6a-g). To further examine whether cystitis-induced changes in DGC*^Trpc4^* neuronal excitability are subtype-specific, we analyzed detailed electrophysiological parameters across the major firing subtypes identified (delayed, phasic, single-spiking, tonic, and gap-firing). To our surprise, we noticed that except the single firing neurons, other *Trpc4*⁺ E-types (delayed, phasic, tonic, and gap-firing) from CYP-treated animals showed significantly altered intrinsic excitability following cystitis (Extended Data Fig.6m), though the magnitude of effect varied across subtypes. This functional plasticity likely contributes to maladaptive processing within lumbosacral spinal circuits underlying bladder pain and voiding dysfunction during cystitis.

## Elevating Endogenous Activity of DGC*^Trpc4^* Interneurons Alleviates Cystitis Phenotypes

Since cystitis attenuates DGC*^Trpc4^* interneuron activity, we tested whether enhancing their excitability could alleviate bladder pain and voiding dysfunction. We used a gain-of-function approach by injecting AAV5-DIO-hM3Dq-mCherry or control AAV5-DIO-mCherry into the lumbosacral DGC of *Trpc4*-Cre mice (Fig.3k-l), enabling chemogenetic activation of DGC*^Trpc4^* interneurons via systemic clozapine-N-oxide (CNO). In naïve mice, activating hM3Dq-expressing DGC*^Trpc4^* interneurons did not alter baseline abdominal sensitivity (Fig. 3m) or nociceptive behavior (Extended Data Fig.7a, b), consistent with prior studies showing spinal interneuron activation has minimal effects without injury ^36–38^, suggesting a state-dependent modulatory role.

To assess whether DGC*^Trpc4^* interneuron activation mitigates cystitis-induced pain, we examined mechanical referred pain and VMRs in cystitis mice. In CYP-treated animals, CNO activation of DGC*^Trpc4^* interneurons significantly reduced abdominal mechanical hypersensitivity (Fig. 3n) and VMRs to bladder distension across pressure ranges (Fig. 3o-p). These effects were absent in Cre-negative controls, indicating that DGC*^Trpc4^* interneuron activation selectively alleviates pain under maladaptive inflammatory conditions without altering baseline sensory processing. We next tested if DGC*^Trpc4^* interneuron activation improves voiding dysfunction in cystitis (Fig. 3q). Cystometric analysis showed that CNO activation in CYP-treated mice significantly lengthened ICIs (Fig. 3r), reducing NVCs (Fig. 3s) and bladder pressure (Extended Data Fig.7c-f), suggesting improved bladder storage and reduced urgency. These changes indicate selective modulation of voiding function in the cystitis state. Collectively, these results position DGC*^Trpc4^* interneurons as a critical regulatory node in lumbosacral circuits capable of restoring bladder homeostasis through bidirectional control of spinal pain and bladder reflex pathways.

## DGC*^Trpc4^* Interneurons Directly Gate Ascending Sensory Transmission

To understand if DGC*^Trpc4^* interneurons directly modulate ascending sensory information flow, we injected retrograde AAV2-tdTomato into the parabrachial nucleus (PBN) and Barrington’s nucleus (BN) and AAV-DIO-ChR2-eYFP into the DGC of *Trpc4*-Cre mice (Extended Data Fig.8a-b). This dual-virus approach enabled selective optogenetic activation of *Trpc4*⁺ axons and electrophysiological recording from spinal projection neurons (SPN) in acute spinal slices. Photostimulation of *Trpc4*⁺ terminals at 0 mV, evoked inhibitory currents that were abolished by TTX and restored by 4-AP, supporting direct GABAergic input from *Trpc4*⁺ neurons (Extended Data Fig.8c-e). Furthermore, at – 70 mV, evoked excitatory currents were abolished by TTX and restored by 4-AP, confirming monosynaptic glutamatergic input (Extended Data Fig.8f–i). These findings indicate that *Trpc4*⁺ DGC neurons provide both excitatory and inhibitory monosynaptic input to SPNs, forming parallel subcircuits capable of bidirectionally modulating ascending visceral sensory transmission, positioning these interneurons as critical gatekeepers of bladder somatosensory signaling within the SC.

## DGC*^Trpc4^* Interneurons Receive Convergent Peripheral and Descending Input

To delineate if DGC*^Trpc4^* interneurons are the gatekeepers of lumbosacral SC sensory processing, we employed a Cre-dependent monosynaptic rabies virus tracing strategy by expressing TVA-GFP, N2C-G protein, and N2C G-deleted rabies in DGC*^Trpc4^*interneurons (Fig. 4a, Extended Data Fig.9a). Consistent with a local integrative role (Extended Data Fig.3, 8), we observed robust labeling of presynaptic neurons within the lumbosacral SC and dorsal root ganglia (DRG). In the DRG, rabies-labeled sensory neurons co-expressed a range of canonical markers, including *NF200* (myelinated Aβ/Aδ fibers), *CGRP* (peptidergic nociceptors), *TH* (C-low threshold mechanoreceptors), *TrkB* and *TrkC* (mechanosensory and proprioceptive afferents), and parvalbumin (PV; proprioceptors) (Extended Data Fig.9b-c). This molecular diversity confirms that DGC*^Trpc4^* interneurons receive direct input from both nociceptive and non-nociceptive primary sensory populations.

**Fig. 4.**
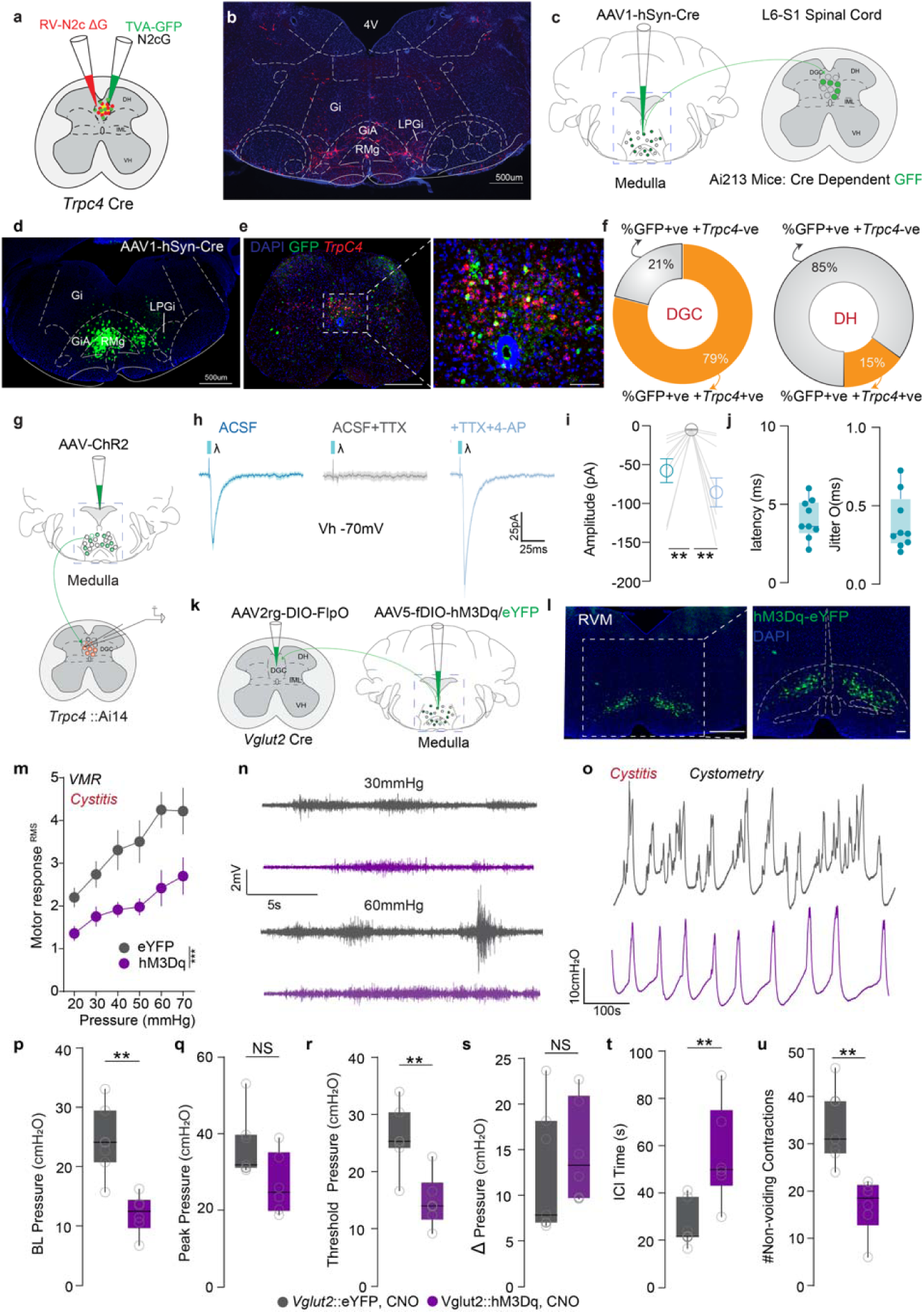
*Vglut2 VMM➔DGC* projections can restore lumbosacral sensorimotor function. (a) Schematic showing monosynaptic rabies tracing from Trpc4⁺ DGC neurons. (b) Representative image reveals retrogradely labeled neurons in the VMM. Scale bar 500um (c) Schematic of transynaptic anterograde tracing using AAV1-hSyn-Cre injection into the VMM of Ai213 mice to label downstream GFP⁺ neurons in the lumbosacral spinal cord. (d) GFP⁺cells in the VMM from AAV1-hSyn-Cre injection. Scale bar 500um. (e) GFP⁺ cells are primarily localized in the DGC and co-express Trpc4, indicating preferential targeting of Trpc4⁺ neurons.(f) Quantification shows that 79% of GFP⁺ DGC neurons are Trpc4⁺, compared to 15% in the dorsal horn.(g) Schematic showing ChR2 expression in the VMM of Trpc4::Ai14 mice.(h–j) Light-evoked EPSCs in tdTomato⁺ Trpc4 neurons are abolished by TTX and rescued by 4AP, confirming monosynaptic inputs.(k) Intersectional strategy targeting Vglut2+ VMMDDGC using AAV2rg-DIO-FlpO and AAV5-fDIO-hM3Dq/eYFP.(l) Representative RVM section showing hM3Dq-eYFP-labeled Vglut2⁺ neurons. Scale bars, 500 µm (overview), 100 µm (zoom). (m) Chemogenetic activation of VMM Vglut2::hM3Dq neurons reduced VMR to bladder distension (***P < 0.001, two-way ANOVA). (n) Representative EMG traces during bladder distensions show decreased VMR in the hM3Dq group. (o) Cystometry recordings reveal improved bladder function following Vglut2+ VMMDDGC neuronal activation. (p–u) Chemogenetic activation Vglut2+ VMMDDGC neurons reduces baseline pressure (p, **P < 0.01, unpaired t test), increases micturition threshold (r, **P < 0.01, unpaired t test), prolongs ICIs (t, **P < 0.01, unpaired t test), and decreases NVCs (U, **P < 0.01, unpaired t test), without affecting peak or Δ pressure (q, s). NS, not significant.

Supraspinally, rabies-labeled neurons were found in the BN, motor cortex (MC), caudal pontine reticular nucleus (PnC), vestibular nuclei, and ventral medial medulla (VMM) (Extended Data Fig.9d-h), implicating descending motor and modulatory inputs. The VMM showed the densest labeling (Fig. 4b), suggesting it is a major source of descending input to DGC*^Trpc4^* neurons. Although BN➔DGC pathways are well characterized in bladder control and appreciated as canonical pathway populations ^39–43^. Roles for other supraspinal inputs, especially the VMM, are less understood. Prior work showed that pseudorabies virus injected into the bladder labels VMM neurons, and that VMM firing tracks micturition episodes ^22,44–48^. These observations suggest that VMM➔SC projections may play an underappreciated role in the descending regulation of bladder function, possibly by modulating spinal interneuron excitability during bladder filling or painful distension.

## *Vglut2***⁺** VMM**➔**DGC Pathway Can Produce Pain Relief and Restores Function in Cystitis

To map this circuit, we injected AAV1-hSyn-Cre into the VMM of Ai213 mice, enabling Cre-dependent GFP expression in downstream spinal neurons (Fig. 4c-d). GFP⁺ neurons were predominantly localized to the lumbosacral DGC, with fewer in the dorsal horn (Fig. 4e). fISH revealed that 79% of GFP⁺ DGC neurons co-expressed *Trpc4*, compared to only 15% in the dorsal horn (Fig. 4f), confirming preferential targeting of DGC*^Trpc4^* neurons by VMM projections. To determine neurotransmitter identity of VMM➔DGC neurons, we performed fISH for *Vglut2* and *Vgat* in rabies-labeled VMM neurons. Over 50% were *Vglut2*⁺, and 41% were *Vgat*⁺ (Extended Data Fig.9i-j), with posterior VMM showing a bias toward excitatory *Vglut2* expression suggesting a rostrocaudal gradient favoring excitatory input (Extended Data Fig.9k). We further confirmed these results by injecting AAV-hSyn-ChR2 into the VMM of *Trpc4*-Cre:Ai14 mice and performing whole-cell recordings from tdTomato⁺ neurons in acute lumbosacral SC slices (Fig. 4g). Brief light pulses evoked both monosynaptic EPSCs and IPSCs in *Trpc4*⁺ neurons. These responses were abolished by TTX and rescued by 4-AP, confirming direct functional monosynaptic connectivity (Fig. 4h-j, Extended Data Fig.9l-n). These findings are consistent with our fISH data showing *Vglut2* and *Vgat* expression in rabies-labeled VMM neurons. Together, these results demonstrate that descending VMM inputs form direct functional synapses onto DGC*^Trpc4^*interneurons, positioning the VMM as a key supraspinal node for modulating spinal circuits involved in bladder sensory gating and pelvic reflex control.

As activation of DGC*^Trpc4^* interneurons alleviated bladder pain and dysfunction, we reasoned if enhancing activity of their upstream excitatory inputs from the VMM could replicate these phenotypes. Using an intersectional viral strategy in *Vglut2*-Cre mice. We selectively targeted VMM *Vglut2*⁺ neurons that project to the lumbosacral DGC. AAV2-retro-DIO-FlpO was injected into the lumbosacral spinal cord, while a Flp-dependent AAV encoding hM3Dq-eYFP (or control eYFP) was injected into the VMM (Fig. 4k-l). In naïve mice, chemogenetic activation of hM3Dq-expressing *Vglut2*⁺ VMM neurons did not alter baseline abdominal sensitivity or nociceptive behavior (Extended Data Fig.10a-b). However, in mice with CYP-induced cystitis, CNO-mediated activation of the *Vglut2*⁺ VMM➔DGC pathway significantly increased abdominal mechanical thresholds compared to eYFP controls (Extended Data Fig.10c), indicating a reversal of referred visceral hypersensitivity. Importantly, responses to somatic stimuli (e.g., paw withdrawal) remained unchanged (Extended Data Fig.10d), suggesting this pathway exerts selective, state-dependent modulation of pelvic sensory processing. Activation of *Vglut2*⁺ VMM➔DGC neurons also significantly reduced VMRs to graded bladder distension in CYP-treated mice (Fig. 4m-n), indicating relief of bladder pain. Cystometric recordings showed decreased bladder pressure and NVCs, along with lengthened ICIs, consistent with improved bladder storage capacity and reduced urgency (Fig. 4o-u). These findings identify the excitatory *Vglut2*⁺ VMM➔DGC pathway as a key modulator of spinal circuits controlling bladder pain and voiding, capable of restoring lumbosacral sensorimotor function in a model of cystitis.

## Discussion

The lumbosacral SC has long been recognized as a crucial neural substrate for integrating afferent signals from the urinary bladder, regulating bladder pain, and coordinating voiding behaviors. Despite this foundational knowledge, the precise identity of the spinal neurons responsible for these functions has remained elusive, limiting our ability to define the underlying circuit logic of bladder somatosensation. Here we identify a genetically defined spinal interneuron population labelled by *Trpc4* expression in the lumbosacral DGC that appears to gate bladder sensory signals and coordinate micturition reflexes. Notably, the DGC is known to receive convergent input from bladder and urethral afferent ^12,14,22^, implying that DGC*^Trpc4^* interneurons sit at a hub of convergent visceral inputs. Single-cell transcriptomics and electrophysiology reveal that *Trpc4*+ DGC neurons are heterogeneous, comprising both excitatory and inhibitory subtypes with stereotyped firing patterns that could coordinate bladder voiding reflex and activity dependent gating of the bladder pain. These findings reveal *Trpc4*+ DGC neurons as gatekeepers of bladder somatosensation, with the capacity to dynamically regulate both pain and voiding reflexes.

Together, these findings reveal that *Trpc4*⁺ neurons in the DGC not only integrate peripheral visceral sensory inputs but also function as a critical hub for top-down modulation from supraspinal centers. This convergence of ascending and descending pathways positions *Trpc4*⁺ neurons as dynamic gatekeepers within spinal circuits that regulate bladder pain and micturition. Importantly, our study also provides a comprehensive single-cell transcriptional atlas of the lumbosacral spinal cord, identifying molecularly distinct *Trpc4*⁺ subtypes and delineating their electrophysiological and morphological properties. In parallel, our circuit-mapping experiments define the input–output architecture of DGC circuits, laying a foundational roadmap for future investigations into how diverse spinal neuron types contribute to the encoding, gating, and execution of bladder-related somatosensory and motor functions.

### Methods Animals

All experimental procedures were approved by the Institutional Animal Care and Use Committee (IACUC) of Washington University School of Medicine and conducted in accordance with the National Institutes of Health guidelines for the care and use of laboratory animals. No more than five animals were housed per cage. C57BL/6J ; strain #000664, Ai14(B6.Cg-Gt(ROSA)26Sortm14(CAG-tdTomato)Hze/J); strain #007914, Ai213(B6;129S6-Igs7tm213(CAG-EGFP,CAG-mOrange2,CAG-mKate2)Tasic/J); strain #034113 and Vglut2-ires-Cre (Slc17a6^tm2Lowl^; strain #028863 were purchased from Jackson Laboratories and colonies were established in our facilities. Trpc4-Cre (Tg(Trpc4-cre)383Stl; MGI:6114650) mouse line obtained from Susumu Tonegawa’s lab ^49^. All transgenic mice were bred onto C57BL/6J background. Patch-clamp was performed on heterozygote Trpc4-Cre mice crossed to homozygous Ai14 or Ai213 mice. For all viral injections and behavioral experiments, male and female mice between 4 weeks and 8 months were randomly assigned to experimental conditions. To reduce animal use, cystometry was performed in male mice and VMR studies were conducted in females. Randomization was performed for the entire study. Experimenters were blinded to both treatment groups and genotypes to avoid bias during the experimental procedures and analysis.

### Viral constructs

Purified and concentrated adeno-associated viruses were used for expressing desired constructs. The viruses included Cre-dependent Casp3 (AAV1-flex-taCasp3-TEVp; UNC), Cre-dependent hM3Dq-mCherry (pAAV-hSyn-hM3D(Gq)-mCherry; Addgene; Catalog# 50474-AAV5; titer ≥ 3×10¹² vg/mL), and control EYFP (AAV5-EF1a-DIO-EYFP; Addgene; Catalog# 27056; titer ≥ 1×10¹³ vg/mL). For Rabies monosynaptic tracing experiments, helper viruses AAV-Ef1α-DIO-H2B-BFP-2a-TVA-WPRE-pA (BrainVTA) and AAV-DIO-Ef1α-FLEX-H2B-GFP-P2A-N2c(G)-WPRE-pA (BrainVTA) were mixed at a 1:1 ratio), N2C G-deleted Rabies-tdTomato (N2c ΔG-tdTomato; Thomas Jefferson University). For electrophysiology experiments, ChR2-eYFP (AAV-EF1a-double floxed-hChR2(H134R)-EYFP-WPRE-HGHpA; Addgene; Catalog# 20298-AAV5; titer ≥1×10¹³ vg/mL), retrograde-labeling tdTomato virus (pAAV-CAG-tdTomato; Addgene; Catalog# 59462AAVrg; titer ≥1.4x10^13) and FlpO (AAVrg pEF1a-DIO-FLPo-WPRE-hGHpA; Addgene; Catalog# 87306-AAVrg; titer ≥1.6×10¹³ vg/mL) and AAV-fDIO-hM3Dq-2A-eYFP (this paper); titer ≥6.75×10^12^ vg/mL were used. All vectors were aliquoted and stored at −80°C until use.

### Surgical Procedures

Stereotaxic viral injections were performed in *Trpc4*-Cre or Trpc4-Cre:Ai14/Ai213 mice to deliver viral vectors into the L6–S1 VMM, or parabrachial nucleus (PBN). Mice were anesthetized with isoflurane (5% for induction; 1.5–2% for maintenance), and the surgical site was shaved, sterilized, and incised along the midline over the target region. Intraspinal Injections: Mice were positioned in a stereotaxic frame (Kopf or RWD) with vertebral clamps secured at the L1 vertebra. A laminectomy was performed to expose the L6–S1 SC by carefully removing the vertebral bone with a microdrill. The posterior spinal artery served as a reference landmark, and the injection site was located 0.1 mm lateral to the vessel on the left side. A pulled glass micropipette (Nanoject, Drummond) was inserted at a 10° angle to a depth of 0.54–0.57 mm (adjusted by mouse age). 100– 150 nL of virus was injected per site at a rate of 60 nL/min, with four injection sites spaced 0.5 mm apart along the L6–S1 axis. After each injection, the micropipette was held in place for 5–10 minutes to allow viral diffusion. Brain Injections: For injections into supraspinal regions, mice were positioned in the stereotaxic frame and the bregma was used as the anatomical reference. The skull was leveled to within ±0.03 mm error. Craniotomies were made at the following coordinates (relative to bregma): VMM: AP −6.2-6.3 mm, ML 0.0 mm, DV −5.8 mm; PBN/BN: AP −4.7-5.0 mm, ML +1.8 mm, DV −4.0 mm, 300–500 nL of virus was injected per site. The micropipette was left in place for 10 minutes post-injection before withdrawal. Animals were monitored continuously until full recovery from anesthesia. For labeling experiments EUS, targeted injections of Cholera Toxin Subunit B (CTB), Alexa Fluor™ 647 Conjugate (Thermo Fisher Scientific, Cat# C34778) were performed.

Mice were anesthetized with isoflurane and placed in the supine position. A midline incision (∼1.5 cm) was made to expose relevant anatomical targets. For access to the EUS, the abdominal musculature was gently retracted using blunt dissection. A sharp glass micropipette or Hamilton syringe was loaded with 2–5 µL of CTB-647 and inserted directly into the target muscle or organ. The tracer was injected at a rate of 500 nL/min using a manual or automated injector. To ensure optimal uptake and diffusion, the pipette or needle was left in place for 10 minutes post-injection before withdrawal. The incision was then closed using 5-0 nylon sutures, and mice were returned to a heated recovery chamber. Animals were monitored daily postoperatively to ensure complete recovery and to assess for any adverse effects. Postoperative care incisions were closed with sutures, and topical triple antibiotic ointment was applied. Mice received analgesics (buprenorphine, 0.05 mg/kg, i.p.) and were placed on a heated recovery pad.

### Visceromotor Reflex Behavior

The visceromotor reflex (VMR) is typically measured by recording the electromyographic (EMG) activities of the abdominal oblique muscles, which is related to visceral nociception. It is a reliable and reproducible measure, performed as previously described ^26,50^. Female mice were anesthetized with 1.5% isoflurane in oxygen for surgical preparation. Two silver wire electrodes were implanted into the external oblique abdominal muscles for EMG recording, and a third wire was inserted subcutaneously as a ground reference. To induce urinary bladder distension (UBD), a lubricated 24-gauge angiocatheter was gently inserted through the urethra into the bladder. Proper catheter placement was verified by the outflow of clear urine, confirming intravesical access without trauma or bleeding. Following instrumentation, anesthesia was gradually reduced from 2% to 0.8– 1% to reach a light surgical plane. Adequacy of anesthesia was confirmed by the presence of a flexion reflex (paw or tail pinch) and absence of the righting reflex, a standard criterion for semi-conscious VMR recording. EMG signals from the abdominal muscles were recorded with an amplifier and data-acquisition software (sampling rate 1000 Hz; LabChart, AD Instruments, Colorado Springs, CO) on a computer system equipped with an analog-to-digital converter (PowerLab, AD Instruments). Baseline EMG activity was recorded prior to bladder stimulation and subtracted from subsequent responses. Phasic UBD was delivered via the angiocatheter in graded pressure steps (20–80 mmHg), increasing in 10 mmHg increments. Each distension trial evoked EMG responses that were rectified, integrated, and normalized to baseline, with the area under the curve (AUC) calculated as the primary outcome metric. This method allowed precise quantification of distension-evoked abdominal muscle contractions, providing a robust index of bladder-related nociceptive processing.

### Cystometry and External Urethral Sphincter (EUS) Electromyography

To assess bladder function and its coordination with external urethral sphincter (EUS) activity, cystometry and simultaneous EUS electromyography (EMG) were performed in anesthetized mice. Mice were initially anesthetized in an induction chamber with 4% isoflurane, and then transferred to a experimental setup equipped with a gas anesthesia mask for maintenance under ∼2% isoflurane. Anesthesia depth was monitored continuously via respiratory rate, pain reflex (toe pinch), and breathing pattern. The lower abdomen was shaved, sterilized with betadine, and a ∼1.5 cm midline incision was made to expose the lower urinary tract (LUT). A 25-gauge needle connected to polyethylene tubing was used to puncture the bladder dome. The catheter was connected to a three-way valve linking an infusion pump (infusion rate: 0.04 mL/min) and a pressure transducer to continuously record intravesical pressure during bladder filling and voiding cycles. Following exposure and catheter placement, urethane anesthesia (1.2 g/kg, i.p.) was administered to ensure stable conditions for bladder physiology recording. Isoflurane was discontinued after urethane delivery. This transition enabled high-fidelity measurement of spontaneous bladder activity without suppression of reflexes.

Simultaneous EUS EMG Recording: To assess sphincter activity in coordination with bladder pressure, the pubic symphysis was cut using spring scissors to expose the EUS. Two fine wire electrodes were inserted percutaneously into the EUS muscle using 30-gauge needles, with each wire tip bent to form a small hook to anchor the electrode within the muscle. EMG signals from the EUS and bladder pressure were recorded with an amplifier and data-acquisition software (sampling rate 1000 Hz; LabChart, AD Instruments, Colorado Springs, CO) on a computer system equipped with an analog-to-digital converter (PowerLab, AD Instruments), enabling precise evaluation of bladder–sphincter coordination during filling, non-voiding contractions, and micturition.

### Mechanical Sensitivity

Abdominal mechanical sensitivity was assessed using calibrated von Frey filaments (North Coast Medical, Inc., Gilroy, CA) applied to the lower abdominal region, as previously described in our earlier work ^26^. Filaments of increasing force (0.02, 0.08, 0.32, 0.64 and 2.56 g) were applied perpendicular to the skin surface until the filament bent slightly, ensuring consistent pressure delivery. Each filament was applied 5 times at 5–10 second intervals, and the number of withdrawal responses (e.g., brisk contraction, licking, or flinching) was recorded. Testing began with the lowest force filament and progressed sequentially to the highest. Mice were habituated to the testing apparatus prior to assessment to minimize stress-related variability. The total number of withdrawals per force level served as a quantitative index of referred mechanical hypersensitivity, particularly in models of visceral pain or inflammation such as cystitis.

For paw mechanical sensitivity, mice were acclimated for at least one hour in individual red-tinted acrylic cylinders enclosures positioned on an elevated mesh platform to allow unobstructed access to the plantar surface of the paws. Each filament was applied 5 times per paw, with a minimum interstimulus interval of 15 seconds and a 5-minute interval between different filament forces. A brisk paw withdrawal upon filament contact was considered a positive response. The number of withdrawals at each force level was recorded and used as a measure of mechanical sensitivity following previously established protocols ^26,51^.

### Thermal Sensitivity (Hargreaves Test)

Thermal nociception was evaluated using previously described ^26,51^. A focused beam of radiant heat (IITC Life Science, Model 390) was applied to the plantar surface of each hind paw, and the paw withdrawal latency was recorded to the nearest 0.1 second. Each paw was tested three times, with the average of six total trials (three per paw) used for analysis. A minimum intertrial interval of 5 minutes was maintained to prevent sensitization or habituation.

### Pinprick

Mechanical sensitivity was assessed using a non-penetrating stimulus applied to the plantar surface of the hindpaw. Mice were placed on an elevated metal grid platform with 2 mm × 2 mm openings and individually housed within red-tinted acrylic cylinders to minimize visual stress and allow clear access to the paws. Animals were acclimated for at least 30 minutes prior to testing. A non-penetrating Austerlitz pin (size 000; Fine Scientific Tools) was gently applied to the plantar surface of each hindpaw in a consistent manner, ensuring that the skin was indented but not pierced. The number of withdrawal responses was recorded across five consecutive trials per paw, completed within 1 minute. A brisk paw withdrawal, flick, or lift was considered a positive response.

### Immunohistochemistry

Mice were deeply anesthetized using a ketamine/xylazine cocktail (dosage per IACUC protocol). Once a surgical level of anesthesia was confirmed by the absence of a paw-pinch reflex, animals were transcardially perfused with 50 mL of ice-cold 1× phosphate-buffered saline (PBS) to flush circulating blood, followed by 50 mL of ice-cold 4% paraformaldehyde (PFA) in PBS for tissue fixation. After perfusion, tissues of interest were dissected, post-fixed overnight in 4% PFA at 4°C, and then cryoprotected by immersion in 30% sucrose in PBS for 16–48 hours at 4°C, until they sank. Samples were embedded in optimal cutting temperature (OCT) compound, frozen on dry ice, and stored at –80°C until sectioning. Frozen sections (15–30 µm thickness) were cut using a cryostat (CM1950, Leica, Germany) and mounted on Superfrost Plus glass slides for immunofluorescence analysis. Slides were washed in 1× PBS, then blocked for 1 hour at room temperature in blocking solution containing 5% normal donkey serum (NDS) and 0.3% Triton X-100 in PBS. Sections were incubated overnight at 4°C with primary antibodies diluted in the same blocking buffer. The following primary antibodies were used:

Anti-RFP (Rabbit, Clontech, Cat# 632543; 1:1000), Anti-GFP (Chicken monoclonal, Aves Labs, Cat# A11122; 1:2000), Anti-*NF200* (Rabbit polyclonal, Sigma-Aldrich, Cat# N4142; 1:500)*, Anti-CGRP* (Mouse monoclonal, Abcam, Cat# ab81887; 1:500), Anti-*TH* (Chicken, Aves Labs, Cat# TYH; 1:1000), Anti-*TrkB* (Goat polyclonal, R&D Systems, Cat# AF1494; 1:500), Anti-*TrkC* (Goat polyclonal, R&D Systems, Cat# AF1404; 1:500), Anti-*PV* (Mouse monoclonal, Synaptic Systems, Cat# 195011; 1:500). After three 5-minute washes in 1× PBS, sections were incubated for 2 hours at room temperature with Alexa Fluor-conjugated secondary antibodies (Life Technologies; donkey anti-chicken/rabbit/sheep IgG, Alexa Fluor 488; 1:500) protected from light. Slides were washed again three times (10 minutes each) in PBS, air-dried in the dark, and cover slipped with DAPI Fluoromount-G (Southern Biotech, USA). Images were acquired using BZ-X800 fluorescence microscope (Keyence) under standardized exposure and filter settings.

### Fluorescence in situ hybridization (fISH)

To preserve RNA integrity for in situ hybridization, mice were transcardially perfused with DEPC-treated, RNase-free 1× PBS, followed by 4% paraformaldehyde (PFA) in DEPC-treated PBS. Tissue was cryoprotected in 30% sucrose, embedded in optimal cutting temperature (OCT) compound, and cryosectioned at 20 μm thickness using a cryostat. Sections were immediately mounted onto Superfrost Plus microscope slides (Fisher Scientific), air-dried for 1 hour at –20 °C, and then stored at –80 °C for up to 3 months. Prior to hybridization, slides were baked at 60 °C for 1 hour in a hybridization oven, then post-fixed in chilled 4% DEPC-treated PFA for 15 minutes at 4 °C. Slides were washed and incubated in hydrogen peroxide for 10 minutes at room temperature to quench endogenous peroxidase activity. Target retrieval was performed by steaming sections (>95 °C) for 5 minutes, followed by Protease III digestion at 40 °C for 30 minutes in a HybEZ™ oven (Advanced Cell Diagnostics). RNAscope® Multiplex Fluorescent V2 Assay (Advanced Cell Diagnostics, ACD) was used to detect target transcripts. Sections were hybridized with the following probes for 2 hours at 40 °C: *Trpc4* (Cat# 591871), Vglut2 (*Slc17a6*), Cat# 319171, Vgat (*Slc32a1*), Cat# 319191. Following probe hybridization, amplification steps (Amp 1–3) were conducted according to the manufacturer’s protocol. Opal fluorophores (Akoya Biosciences; Opal 520, 570, and 650) were used to visualize distinct transcripts through separate fluorescence channels. Nuclei were counterstained with DAPI, and slides were cover slipped using an antifade mounting medium. Images were acquired using BZ-X800 fluorescence microscope (Keyence)

### Acute SC slice preparation

On the experimental day, mice were anesthetized with ketamine (100 mg/kg) and xylazine (10 mg/kg). Following confirmation of surgical anesthesia by negative tail pinch response, transcardial perfusion was performed using ice-cold, oxygenated N-methyl-D-glucamine (NMDG) solution containing: 93 mM NMDG, 2.5 mM KCl, 1.25 mM NaH₂PO₄, 30 mM NaHCO₃, 20 mM HEPES, 25 mM glucose, 5 mM ascorbic acid, 2 mM thiourea, 3 mM sodium pyruvate, 10 mM MgSO₄, 0.5 mM CaCl₂, and 12 mM N-acetylcysteine (pH 7.3, adjusted with 12 N HCl). The lumbosacral SC segment was rapidly extracted and extruded using a syringe filled with chilled NMDG solution. For virus-injected animals, manual SC dissection was performed to prevent potential post-surgical tissue adhesions. Transverse sections (250–300 μm) were prepared using a Compresstome vibratome (VF-510-0Z, Precisionary Instruments, USA) in ice-cold NMDG solution. Slices were allowed to recover in pre-warmed, oxygenated NMDG solution at 35°C for 10 minutes, then transferred to oxygenated (95% O₂–5% CO₂) artificial cerebrospinal fluid (aCSF) containing: 124 mM NaCl, 2.5 mM KCl, 1.25 mM NaH₂PO₄, 24 mM NaHCO₃, 5 mM HEPES, 12.5 mM glucose, 1 mM MgCl₂, and 2 mM CaCl₂ (pH 7.3, adjusted with 10 M NaOH). Slices were maintained at room temperature for a minimum of 30 minutes before recording.

### Whole-cell slice electrophysiology

Patch-clamp recording experiments were performed at 34°C using a heated recording chamber (ALA Scientific Instruments, HCS) and temperature controller (ALA, HCT-10) in slices obtained as above. Spinal DGC neurons were identified in the recording chamber of a SliceScope Pro 6000 (Scientifica) under a 40X objective (0.8 numerical aperture, Olympus), using an infrared DIC camera (Electro, Teledyne Photometrics). Recording pipettes were fabricated from borosilicate glass capillaries (BF150-110-10HP, Sutter Instrument, USA) using a P-1000 micropipette puller (Sutter Instrument, USA), yielding tip resistances of 4–8 MΩ. Data acquisition was performed using an Axon Instruments DigiData 1550B digitizer, MultiClamp 700B amplifier, and pClamp 11 software (Molecular Devices, USA) at a sampling rate of 2 kHz. For firing pattern characterization, a potassium gluconate-based internal solution was used containing: 120 mM K-gluconate, 5 mM NaCl, 2 mM MgCl₂, 0.1 mM CaCl₂, 4 mM Na₂ATP, 0.4 mM Na₂GTP, 15 mM Na₂-phosphocreatine, 1.1 mM EGTA, and 10 mM HEPES (pH 7.3, adjusted with 10 M KOH; 290 mOsm). Current-clamp recordings were obtained at resting membrane potential using depolarizing current steps ranging from -50 pA to +90 pA in 10 pA increments, with each step lasting 10 seconds. Action potential firing patterns were classified into five categories based on responses to depolarizing current injections that evoked maximal spike numbers: (1) Single spike neurons fired only one action potential regardless of current amplitude; (2) Delayed firing neurons exhibited first action potentials with consistent delays >100 ms; (3) Gap firing patterns were characterized by distinct silent periods; (4) Phasic firing neurons produced two or more bursts of action potentials separated by silent periods ≥100 ms; (5) Tonic firing neurons generated multiple action potentials at threshold depolarizations, with stronger current injections producing high-frequency firing (up to 50 Hz) often accompanied by frequency adaptation. For postsynaptic current recordings, the potassium gluconate internal solution was replaced with a cesium gluconate-based solution containing: 110 mM D-gluconic acid, 110 mM CsOH, 0.1 mM CaCl₂, 4 mM MgATP, 0.4 mM Na₂GTP, 10 mM Na₂-phosphocreatine, 1.1 mM EGTA, 10 mM HEPES, and 8 mM TEA-Cl. Light-evoked postsynaptic currents (eEPSCs or eIPSCs) were recorded at holding potentials of -70 mV or 0 mV in the aCSF to determine the presence of monosynaptic inputs. After obtaining a whole-cell recording, ChR2 was stimulated by flashing (5 ms) 473 nm light through the light path of the microscope using an LED illumination (Thorlabs, USA) by a LED driver. To confirm the polysynaptic or monosynaptic connections, evoked-EPSC or IPSCs were recorded in the aCSF and followed by adding the 1 μM tetrodotoxin (Hello Bio, USA) to aCSF, then a combination of TTX (1μM) and 4-AP (100 μM).

### Morphology reconstruction

For morphological analysis of recorded neurons, the internal solution was supplemented with 0.25% biocytin during patch-clamp recordings. Following completion of electrophysiological recordings, the recording pipette was carefully withdrawn from the patched cell, and the slice was returned to the incubation chamber for a minimum of 30 minutes to allow biocytin diffusion throughout the cell. Slices were subsequently fixed in 4% paraformaldehyde (PFA) in 1× phosphate-buffered saline (PBS) at 4°C for 1–2 overnight periods. Following fixation, slices were washed three times with 1× PBS and permeabilized with 0.3% Triton X-100 in 1× PBS. Biocytin-filled neurons were visualized using Alexa Fluor 647-conjugated streptavidin (1:500 dilution; Thermo Fisher Scientific, USA) applied overnight at 4°C. After three additional washes with 1× PBS, slices were mounted using DAPI Fluoromount-G mounting medium (Southern Biotech, USA). Fluorescent images were acquired using a DMI4000 B confocal microscope (Leica, Germany). Morphological reconstruction (skeletonization) and Sholl analysis of neurite intersections were performed using the Simple Neurite Plugin ^52^ of Fiji ImageJ.

### Electrophysiology feature analysis

To extract and analyze electrophysiological features from whole-cell recordings, we used a custom Python pipeline leveraging the pyABF package and the *Electrophys Feature Extraction Library* (eFEL) to load and process .abf files. Recordings were organized by folder, with each folder representing a distinct firing pattern group (e.g., *Gap*, *Single*, etc.). Extracted features were standardized across the dataset using z-score normalization. To determine which features were significantly associated with firing pattern groups, we performed both one-way ANOVA and Kruskal–Wallis tests for each feature. Only parameters with a p-value < 0.05 in both test were retained for subsequent analysis, resulting in a filtered feature matrix. All statistical tests were performed using the SciPy package, and data manipulation was conducted in NumPy and Pandas DataFrame format. To visualize group-specific feature distributions, we generated a clustered heatmap of the filtered features comparing five firing pattern groups. Hierarchical clustering was applied to the features (columns) to group similar parameters together, enhancing interpretability of the heatmap. For treatment comparisons, the dataset was further stratified by saline or CYP treatment within each firing pattern group. Within-group one-way ANOVA was used to identify features that significantly differed between treatment conditions (p < 0.05). These differentially expressed features were pooled across all firing types and used to construct a second heatmap, illustrating treatment-dependent changes in electrophysiological parameters across firing pattern subtypes. Finally, to explore high-dimensional structure in the dataset, we applied t-distributed Stochastic Neighbor Embedding (t-SNE) using the *scikit-learn* library. The filtered and normalized feature set was projected into two dimensions, and the resulting scatter plot was color-coded by firing pattern group to visualize separability in reduced feature space. All plots generated in this pipeline use matplotlib and seaborn.

### Patch-Seq

To molecularly profile electrophysiologically characterized neurons, we performed patch-seq on a subset of tdTomato+/GFP+ neurons recorded in the lumbosacral DGC of Trpc4:Ai14/Ai213 mice. Slice preparation and recording conditions were matched to those used for whole-cell patch-clamp electrophysiology performed above. The internal pipette solution was modified from standard composition, based on previously published ^53^ patch-seq protocol, to optimize RNA preservation and cDNA amplification. The solution contained (in mM): 110 potassium gluconate, 10 HEPES, 0.2 EGTA, 4 KCl, 0.3 GTP (sodium salt), 10 phosphocreatine (disodium salt), 1 ATP (sodium salt), and supplemented with 20 µg/mL glycogen, 0.2 U/µL RNase inhibitor (Roche, # 03335402001), and 3mg/ml biocytin (Sigma, B4261), adjusted to pH 7.4 with KOH. Fill the pipette with 1.8ul internal solution. After completion of electrophysiological characterization, the recording pipette was carefully positioned over the soma or nucleus (if visible). Mild negative pressure (∼–30 mbar) was applied to extract the cytosol and attract the nucleus to the pipette tip. After ∼1 minute of gentle aspiration, the soma visibly collapsed and/or the nucleus approached the pipette tip. While maintaining negative pressure, the pipette was slowly retracted in the x and z axes under continuous visual monitoring. The pipette was then removed from the headstage, gently fracture the pipette tips which contain internal solution, cytosolic RNA, and nucleus into a PCR tube pre-loaded with lysis buffer (Takara, #634894). Samples were flash-frozen and stored at –80°C until subsequent library preparation and sequencing. Following nucleus extraction, single-cell lysates were processed using the SMART-Seq v4 Ultra Low Input RNA Kit for Sequencing (Takara, #634894) according to the manufacturer’s protocol. RNA was reverse transcribed and full-length cDNA was amplified with 20 PCR cycles. Each amplification batch included four no-template (H2O) controls and four controls of 50 pg total RNA. Amplified cDNAs were assayed using an Agilent TapeStation 4200. Samples containing more than 1 ng total cDNA or at least 25% of cDNA in the size range of 500 – 6000 bp were deemed high quality and subjected to library preparation using the Nextera XT DNA Library Prep Kit (Illumina, FC-131-1096), incorporating Nextera XT Index Kit v2 Sets A–D (FC-131–2001, 2002, 2003, 2004). Sequencing libraries were pooled and sequenced on an Illumina NovaSeq X Plus platform, targeting a sequencing depth of at least 200k reads per sample. Sequencing reads were aligned to mm10 Reference genome and gene expression counts were generated.

### Patch-seq data analysis

Gene expression data of Patch-seq samples were loaded into R package Seurat (V5.1.0) for processing. To control for sample quality, Patch-seq samples with less than 1,000 genes were excluded from the analysis. Gene expression counts were normalized using the NormalizeData() function to account for differences in sequencing depth. Data were subsequently scaled and centered using the ScaleData() function. To identify genes that are differentially expressed across firing patterns identified based on electrophysiological recording, differential expression analysis was conducted using the FindAllMarkers() function with algorithm set to DESeq2, comparing samples from a given firing pattern to all other samples. Genes with FDR < 0.05 were reported.

### Tissue Collection and Single-Nuclei Isolation

To collect mouse lumbosacral SC tissues for sequencing, animals were anesthetized with a ketamine/xylazine cocktail and transcardially perfused with ice-cold NMDG-based cutting solution described above. The SC was rapidly removed and dissected while submerged in ice-cold NMDG-based cutting solution. SC slices (400 µm thick) were obtained using a Compresstome (Precisionary Instruments, VF-210-0Z). Tissues were stored at -80C until use. Frozen tissue samples were transferred from −80 °C and briefly thawed on ice for 2 minutes. Tissues were transferred to a Dounce homogenizer (Millipore Sigma, D8938-1SET) containing ice-cold homogenization buffer composed of: 0.25 M sucrose, 25 mM KCl, 5 mM MgCl₂, 10 mM Tris-HCl (pH 8.0), 5 μg/mL actinomycin D, 1% bovine serum albumin (BSA), 0.08 U/μL RNase inhibitor, and 0.01% NP-40. Homogenization was performed using 10 strokes with the loose pestle followed by 10 strokes with the tight pestle in a total volume of 1 mL. The resulting homogenate was filtered through a 50 μm mesh filter and centrifuged at 500 × g for 10 minutes at 4°C. Pellets were resuspended in resuspension buffer supplemented with 5 ng/μL 7-aminoactinomycin D (7-AAD). To eliminate cellular debris, nuclei were subjected to fluorescence-activated cell sorting (FACS) using a BD FACSAria III cell sorter equipped with a 100 μm nozzle. 7-AAD-positive events were sorted and collected into sterile 1.5 mL microcentrifuge tubes.

### snRNA-seq

Sorted nuclei were quantified using a hemocytometer and diluted to the appropriate concentration to target recovery of 10,000 nuclei per sample. Single-nucleus RNA sequencing (snRNA-seq) libraries were prepared using the 10× Genomics Chromium Single Cell 3’ V4 GEM-X kit or Multiome kit following the manufacturer’s instructions. Libraries were sequenced on either Illumina NovaSeq 6000 or NovaSeq X Plus using 300-cycle paired-end sequencing (150 cycles for both Read 1 and Read 2), targeting a sequencing depth of 50,000 paired-end reads per nucleus. Sequencing reads were mapped to mouse mm10 reference genome and genes counts were generated using cellranger (version 9.0.1 for GEM-X, 10X Genomics) or cellranger-arc (version 2.0.1 for Multiome, 10X Genomics).

### Quality Control and Clustering of snRNA-seq Data

Single-nucleus RNA sequencing (snRNA-seq) data were analyzed using the Seurat R package (V5.1.0),. Quality control filtering was applied to exclude low-quality nuclei minimum of 500 detected genes per nucleus, fewer than 15,000 total unique molecular identifiers (UMIs), and less than 2% mitochondrial gene counts. Multiplets were identified using DouletFinder and subsequently excluded form the analysis. Gene expression counts were normalized using the NormalizeData() function, which scales each nucleus to 10,000 total transcripts to account for differences in sequencing depth. Data were subsequently scaled and centered using the ScaleData() function. Highly variable genes were identified using FindVariableFeatures(), and dimensionality reduction was performed via principal component analysis (PCA) using RunPCA(), retaining the top 20 principal components (PCs). Uniform Manifold Approximation and Projection (UMAP) was employed for data visualization using RunUMAP() based on the top 20 PCs. Unsupervised clustering was performed using the FindClusters() function with a resolution parameter of 0.6. Marker genes for each cluster were identified using FindAllMarkers() by comparing each cluster against all others, applying significance thresholds of false discovery rate (FDR) < 0.05 and log₂ fold-change > 1. Clusters were flagged as low-quality or potential doublets based on two criteria: (1) absence of significantly enriched marker genes (FDR < 0.05, log₂FC > 1), (2) presence of five or more mitochondrial genes among the top 20 markers (ranked by average log₂ fold-change), or (3) Highly express marker genes (genes see below, FDR < 0.05, log₂FC > 1) for multiple cell types. Cell type annotation was performed based on expression of canonical markers: neuronal clusters were identified by high expression of neuronal markers (e.g., *Rbfox3*, *Snap25*, *Syt1*), while non-neuronal clusters expressed glial or other cell type markers (e.g., *Sparc*, *Mbp*, *Apoe*). Next, we classified each neuronal cluster into the “glutamatergic”, “GABAergic” and “cholinergic” based on their neurotransmitter expression. Clusters were annotated as “glutamatergic” if the expression of *Slc17a6* was higher; clusters were annotated as “GABAergic” if expression of the expression of *Slc32a1* was higher; and clusters were annotated as “cholinergic” if the expression of *Chat* was higher. Initial clusters containing nuclei of multiple neuronal classes were sub-clustered and classified according to the process mentioned above.

### Differential gene expression analyses

To identify transcriptional features, differential expression analyses were conducted using the FindAllMarkers() function in Seurat, comparing nuclei from a given cell type to all other nuclei. Genes with FDR < 0.05 were considered significant. For pairwise comparisons between two specific populations, FindMarkers() was used with FDR < 0.05.

### Integration of published scRNA-seq SC data

snRNA-seq data of mouse lumbosacral SC and DGC neurons generated in this study were integrated with the published single cell data of mouse lumbar SC neurons using Seurat Seurat CCA ^21^. The published dataset was then annotated by transferring the cell type labels reported in this study to individual nuclei of the published dataset using FindTransferAnchors() and TransferData().

### Statistical Analysis

All data analyses were conducted with experimenters blinded to the experimental conditions. Exclusion criteria included failure to localize expression in the experimental model or off-target virus/drug delivery. At least three replicates per condition were performed, and data were averaged for each behavioral assay. The number of animals used in each experiment is denoted by “N.” For statistical comparisons, normality was first assessed using the D’Agostino and Pearson omnibus normality test. Parametric tests, including paired t-tests and two-way ANOVA with Bonferroni post hoc corrections, were used when normality assumptions were met. If normality was not assumed, nonparametric tests (e.g., Wilcoxon matched-pairs test, Mann-Whitney U test) were applied. Statistical significance was set at p < 0.05 for all tests (Extended Data Table 2 shows the values).

## Data and code availability

Raw and processed data of snRNA-seq experiments included in this study are deposited to the NCBI Gene Expression (GEO) SRA with accession number GSEXXXXX. All analysis codes can be downloaded from https://github.com/Samineni-Lab.

## NIH Rights Statement

This manuscript is the result of funding in whole or in part by the National Institutes of Health (NIH). It is subject to the NIH Public Access Policy. Through acceptance of this federal funding, NIH has been given a right to make this manuscript publicly available in PubMed Central upon the Official Date of Publication, as defined by NIH.

## Acknowledgements

This work was funded by the Urology Care Foundation Research Scholars Award to YZ. Takeda Science Foundation Postdoctoral Scholarship to MK. NIDDK Career Development award (K01 DK115634), NIDDK R01DK128475, R01DK139386, R01DK140445 to VKS. We thank the Flow Cytometry & Fluorescence Activated Cell Sorting Core in the Department of Pathology and Immunology at Washington University School of Medicine for help with FACS. We thank the Genome Technology Access Center at McDonnell Genome Institute at Washington University School of Medicine for help with snRNA-seq library construction and sequencing. The Center is partially supported by NCI Cancer Center Support Grant #P30 CA91842 to the Siteman Cancer Center from the National Center for Research Resources (NCRR), a component of the National Institutes of Health (NIH), and NIH Roadmap for Medical Research. We thank Hugo Greenhill and Megha Jacob for help with manuscript preparation. Robert Gereau IV for advice with data interpretation.

## Author contributions

Y.Z., F.L., and V.K.S. designed the experiments. Y.Z. and F.L. performed the surgeries. Y.Z., F.L., H.J.H., M.H., and V.K.S. conducted the behavioral experiments. F.L., H.J.H., Y.Z., N.H., and E.C.MR. performed the anatomical analyses. Y.Z. and L.Y. prepared the libraries and performed FACS for RNA-seq. L.Y., A.P conducted data curation, analysis, and developed methodology for RNA-seq. H.A.H. developed code for cystometry and cell counting. Y.Z., T.O. performed slice electrophysiology and morphology reconstruction. Y.Z., T.O., Z.L., and M.K. analyzed the electrophysiology data. Y.Z., F.L., L.Y., M.K., and V.K.S. wrote the manuscript with input from all authors.

## Extended Data figure legends

**Extended Data Fig. 1.**
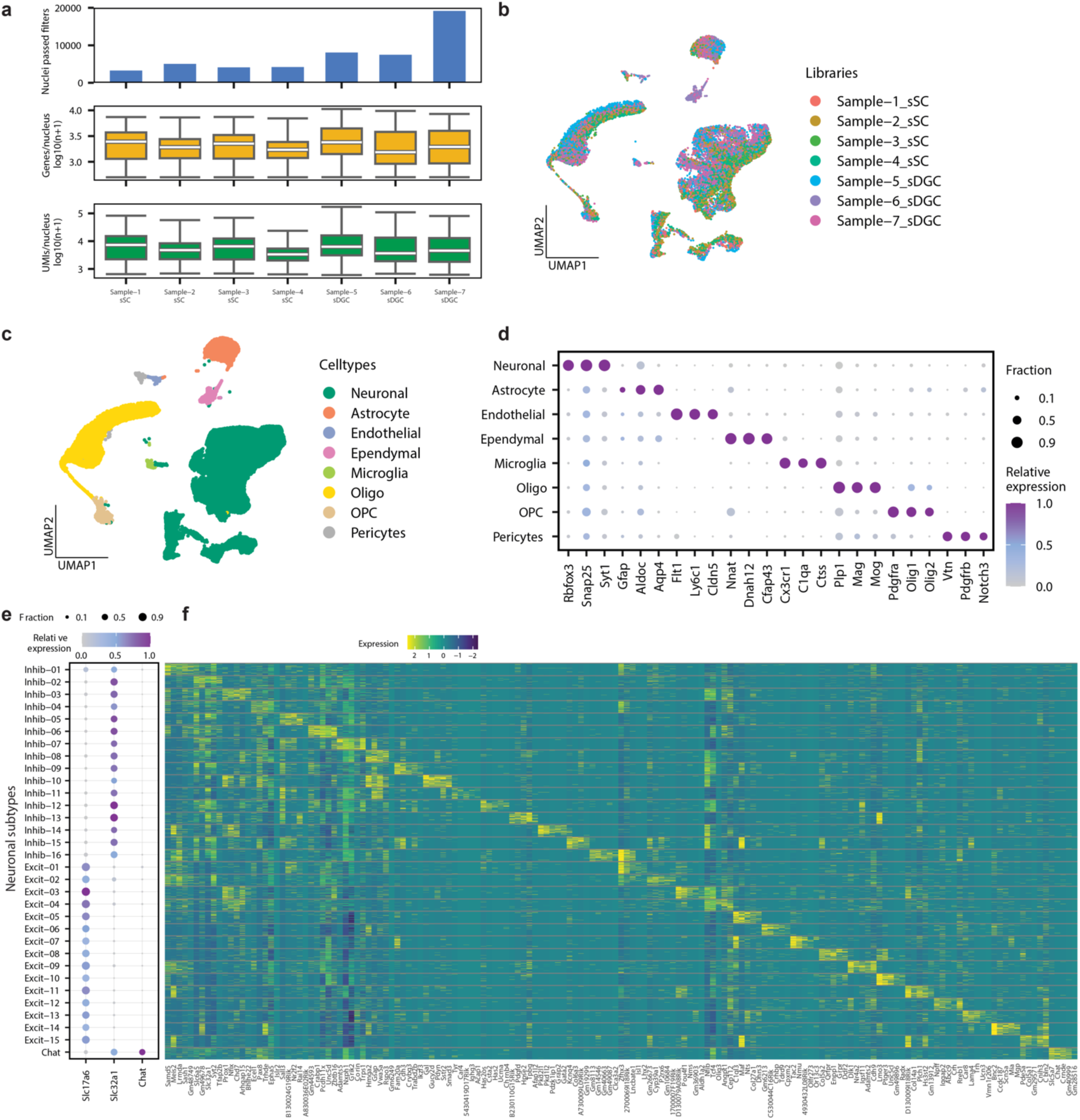
Quality control, cell type identification and marker gene expression in lumbosacral SC. (a) snRNA-seq library metrics. The Top row displays number of nuclei passed quality control. The Middle row displays box plots of number of genes per nucleus (log10 transformed) ,and the bottom row displays the number of UMIs per nucleus. Boxes indicate quartiles and whiskers are 1.5-times the interquartile range (Q1-Q3). The median is a white line inside each box. The distribution is aggregated across all samples and displayed on the horizontal histogram. (b) UMAP of 2,000 downsampled nuclei per library. Nuclei were colored by libraries. (c) UMAP of all snRNA-seq nuclei, colored by cell types. (d). Dot plots showing the marker genes used to annotate each cell type. (e) Dot plots showing the neurotransmitter genes used to classify each neuronal subtype. (f) Heat map showing the expression of top five genes (sorted by Log2FC and FDR) expressed in each neuronal subtype.

**Extended Data Fig. 2.**
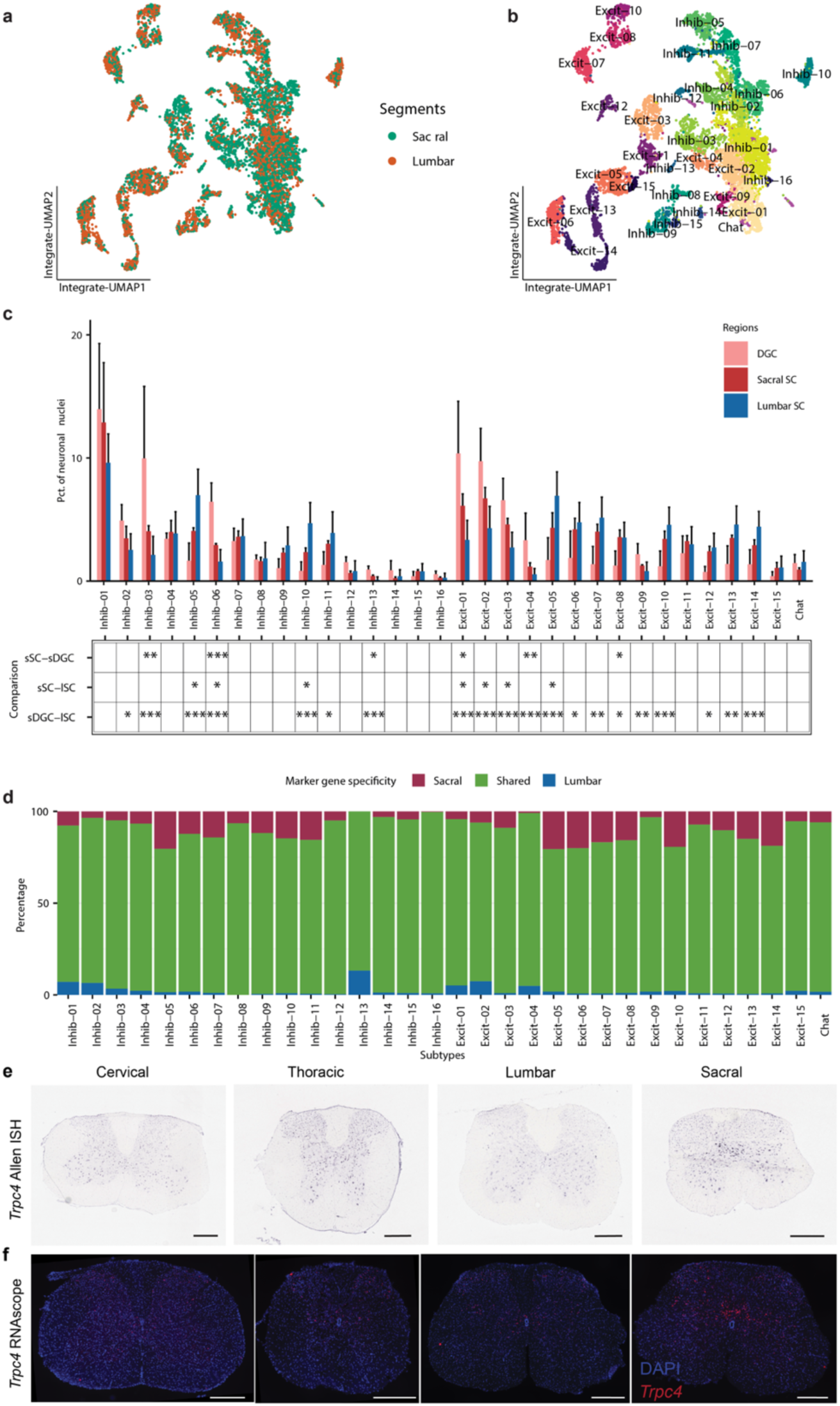
Comparative single-cell analysis of cell type diversity between lumbar and sacral SC. (a) snRNA-seq UMAP showing the integrative analysis of neuronal nuclei from sacral SC and DGC reported in this study and those from lumbar SC previously reported. Nuclei are colored by datasets. (b) snRNA-seq UMAP showing the neuronal nuclei from lumbar SC, sacral SC, and DGC. Nuclei are colored by neuronal subtypes. (c) Bar plot showing the percentage of each neuronal subtype in snRNA-seq libraries separated by regions. Table below shows the significance level of Post-hoc Tukey’s HSD following ANOVA. *p<0.05, **p<0.01, ***p<0.001. (d) Bar plot showing the distribution of neuronal subtype-specific marker genes based on expression between lumbar and sacral SC. Differential expression analysis was done comparing nuclei from sacral SC of one neuronal subtype to nuclei of the same neuronal subtype from lumbar SC. Genes with Log2FC>1 and FDR<0.05 were classified as higher in sacral, genes with Log2FC<(-1) and FDR<0.05 were classified as higher in lumbar. All other genes are classified as shared. (e) ISH images from Allen Brain spinal atlas showing expression of *Trpc4* expression in all segments of SC. (f) RNAScope showing the localization of *Trpc4* in all segments of SC.

**Extended Data Fig. 3.**
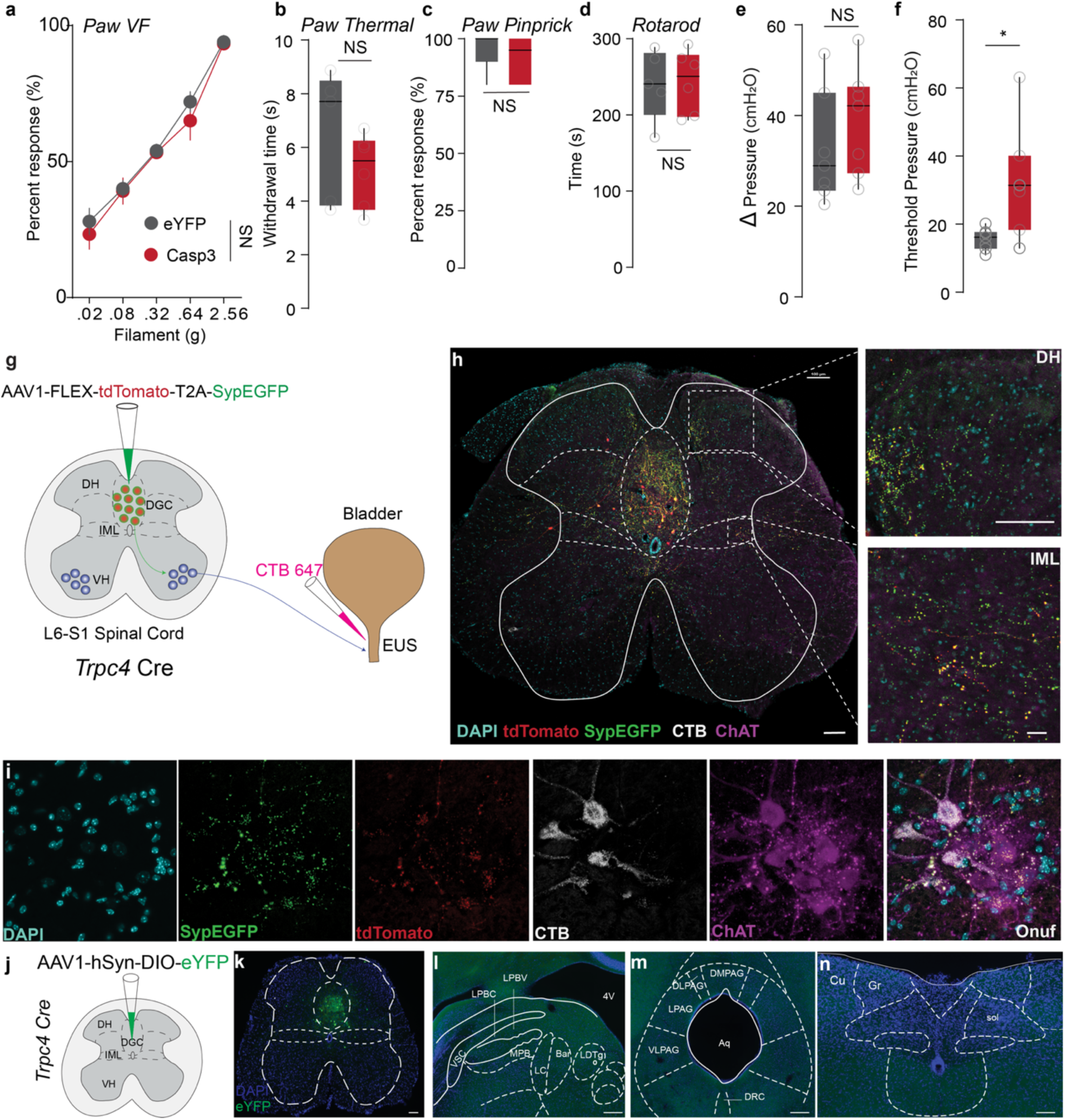
Ablation of DGC*^Trpc4^* local interneurons does not impair somatic or motor behavior. (a) Paw withdrawal responses to von Frey stimulation show no difference between eYFP and Casp3 mice (P = 0.3213, two-way ANOVA). (b–c) Paw withdrawal latency to thermal stimuli (b, P = 0.2867, unpaired t test) and response rate to pinprick (c, P = 0.5455, Mann–Whitney U test) are also unchanged. (d) Rotarod performance time is unaffected by *Trpc4*⁺ neuron ablation (P = 0.9203, unpaired t test). (e) Bladder pressure change (Δ pressure = peak – baseline) during cystometry is comparable between groups (P = 0.3503, unpaired t test). (f) However, the threshold pressure required to elicit micturition is significantly lower in Casp3 mice (*P < 0.05, unpaired t test), suggesting increased bladder sensitivity. (g) Schematic of AAV1-hSyn-DIO-eYFP injection into the lumbosacral dorsal gray commissure (DGC) of *Trpc4*::Cre mice. (h) Representative spinal cord section confirms eYFP-labeled *Trpc4*⁺ neurons are confined to the DGC. (i–k) Examination of brainstem regions, including the parabrachial complex (i), periaqueductal gray (j), and nucleus of the solitary tract, cuneate, and gracile nuclei. (k), No eYFP⁺ axonal projections, indicating that DGC*^Trpc4^* neurons are local spinal interneurons lacking long-range supraspinal connectivity. Scale bars, 100 µm. NS, not significant.

**Extended Data Fig. 4.**
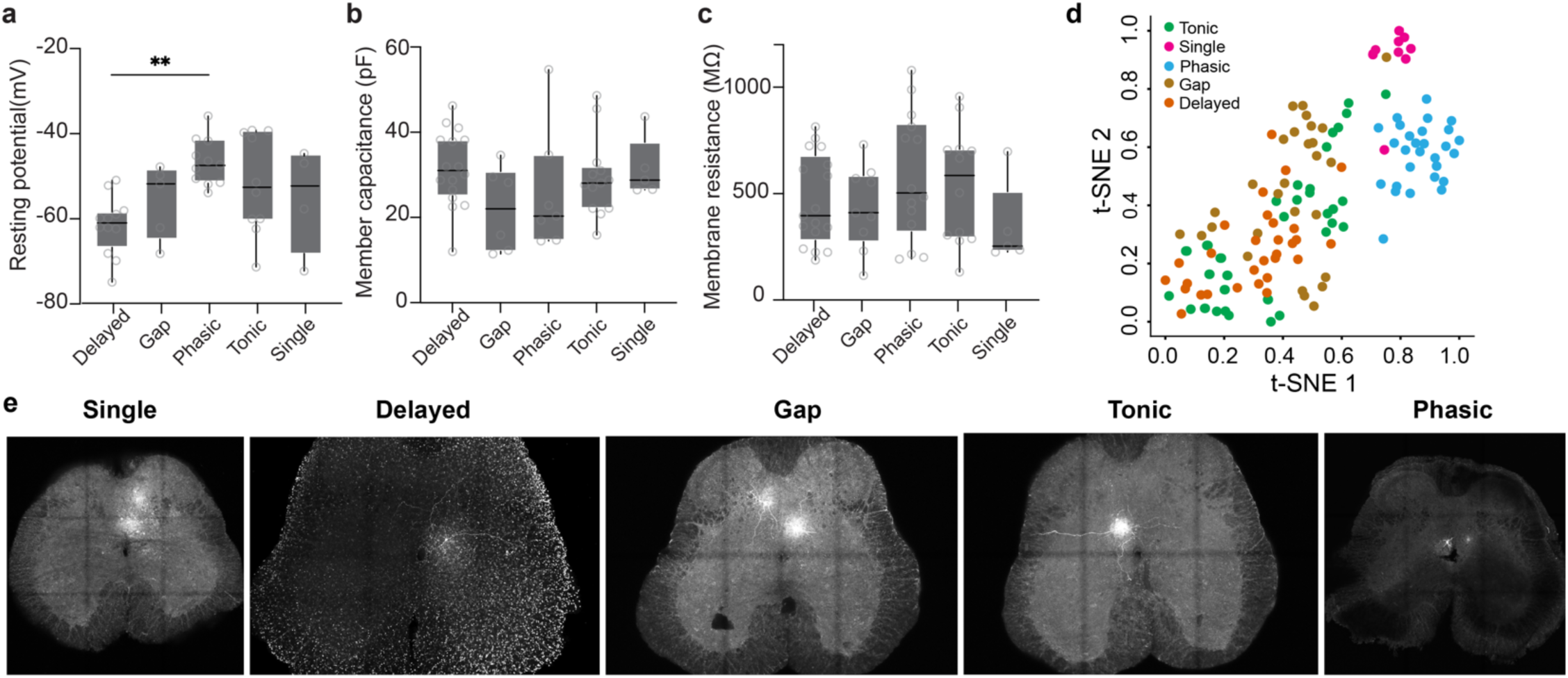
Functional and morphological diversity of DGC*^Trpc4^* neurons. (a–c) Intrinsic membrane properties of identified E-types of *Trpc4*⁺ neurons. (a) Delayed neurons show a significantly more hyperpolarized resting membrane potential (**P < 0.01, one-way ANOVA), while membrane capacitance (b) and resistance (c) are comparable across E-types. (d) t-SNE analysis shows clustering of cells into two distinct subtypes after integrating morphology and E-types of all recorded neurons. (e) Representative spinal cord images of biocytin-labeled neurons illustrating their soma location, dendritic arbors, and axonal projections within the lumbosacral SC for each E-type.

**Extended Data Fig. 5.**
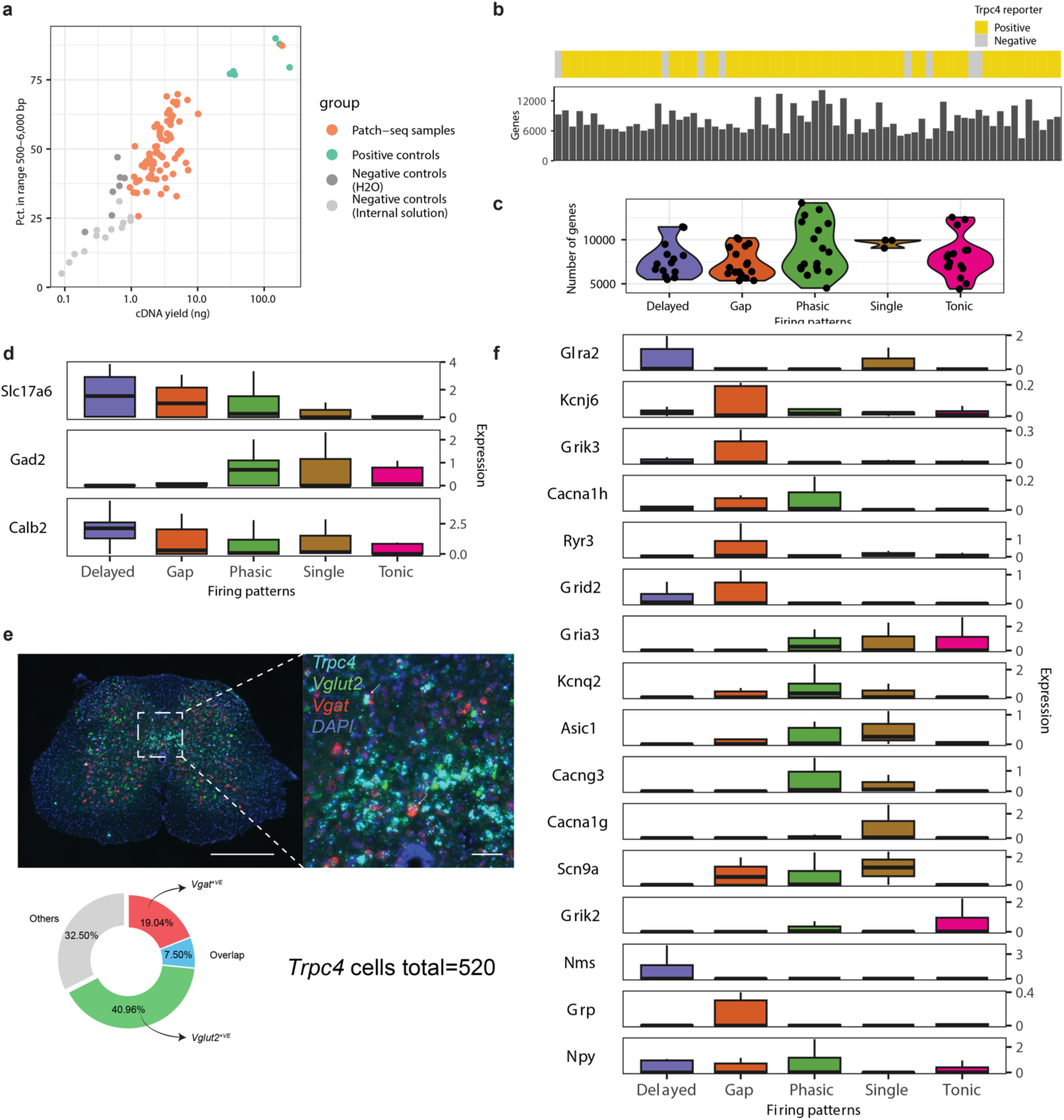
Patch-seq analysis of E-types. (a) Scatter plot showing the cDNA quality control metrics of Patch-seq samples. Samples with at least 2 ng cDNA yield and at least 25% cDNA falling in the range 500-6000 bp were subjected to sequencing and included in the analysis. (b) Top: The expression of Trpc4 reporter for each Patch-seq sample based on their fluorescent properties during Patch-clamp recording. Bottom: bar plot showing the number of genes detected in each Patch-seq samples. (c) Violin plot showing the number of genes detected in samples grouped by their firing patterns (p=0.18, one-way ANOVA). (d) Bar plot showing the expression of selected differentially expressed genes (FDR<0.05) across samples of different firing patterns. (e) RNAScope showing the colocalization of *Trpc4*, *Vgut2*, and *Vgat* in DGC cells. (f) Violin plot showing the expression of previously reported SC interneuron marker genes in Patch-seq samples. Samples are colored by their firing patterns.

**Extended Data Fig. 6.**
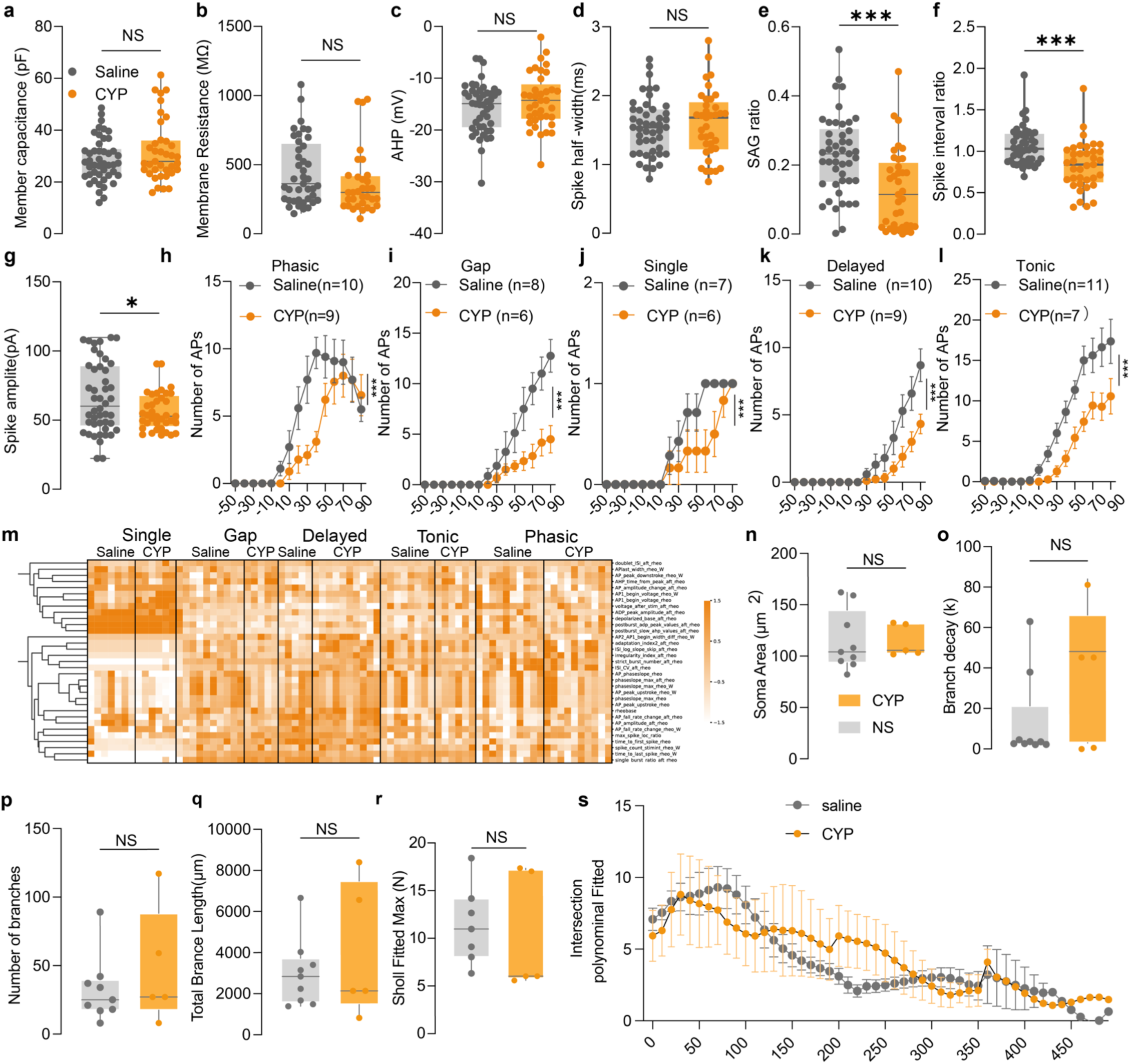
Physiological and morphological classification of DGC*^Trpc4^*neurons following cystitis. (a–f) Summary of (a) membrane capacitance, (b) resistance, (c afterhyperpolarization (AHP), and spike half-width (d) shows no significant difference between saline and CYP-treated groups. While the SAG ratio (e) significantly decreased in CYP-treated neurons (***P < 0.001, unpaired t test). (f) Action potential interval ratio is significantly elevated in CYP group (***P < 0.001, unpaired t test). (g) Spike amplitude is significantly reduced in the CYP group (*P < 0.05, unpaired t test). (h–l) Cystitis significantly reduces DGC*^Trpc4^* neuron excitability, evidenced by fewer action potentials across all neuronal subtypes. (m) Heatmap summarizing electrophysiological parameters analyzed across subtypes and conditions. (n–s) Soma size (n) and dendritic branch decay (o), number of branches (p), total branch length (q), Sholl maximum intersection (r), and polynomial fitting curves from Sholl analysis (s) indicate no significant structural remodeling of DGC*^Trpc4^*neurons in CYP mice compared to saline controls. NS, not significant.

**Extended Data Fig. 7.**
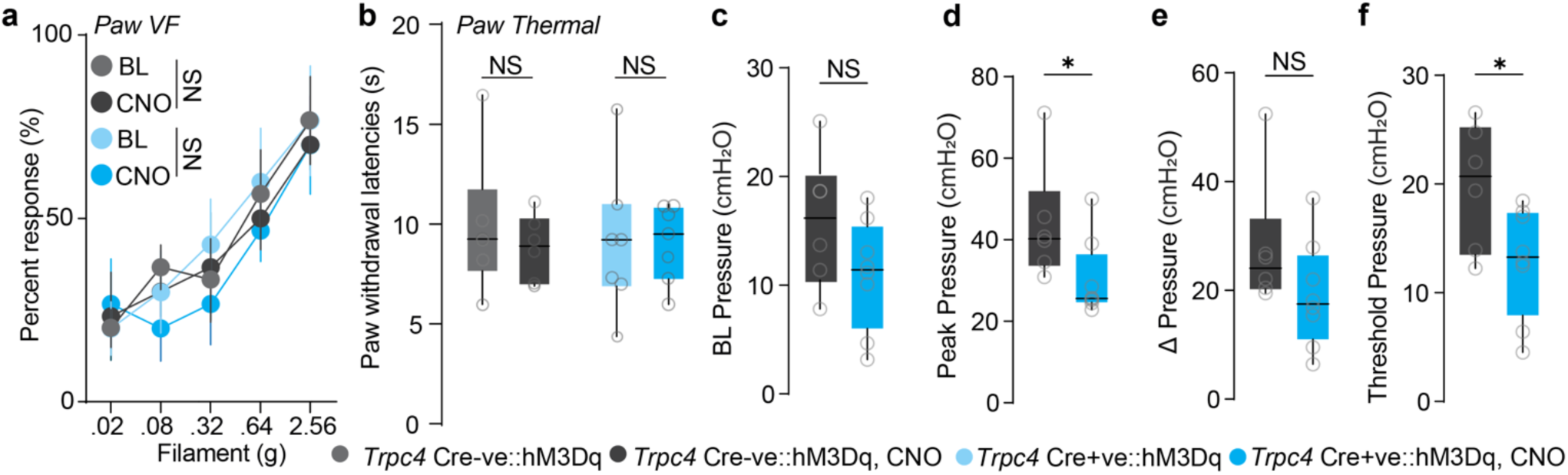
Behavioral effects of activation of DGC*^Trpc4^* neurons. (a) Percent withdrawal response to graded paw von Frey stimulation before (BL) and after CNO injection in *Trpc4*::Cre⁻ and Cre⁺ cystitis mice shows no significant change (P = 0.5560, two-way ANOVA). (b) Paw withdrawal latency to thermal stimulation is also unchanged (P = 0.5111 and P = 0.9516, unpaired t test). (c) Baseline bladder pressure does not differ between groups (P = 0.1309, unpaired t test). (d) Peak bladder pressure is significantly decreased in *Trpc4*::Cre⁺ cystitis mice after CNO injection (*P < 0.05, Mann–Whitney U test). (e) Δ pressure remains unchanged (P = 0.1419, Mann–Whitney U test). (f) Bladder distension threshold pressure is decreased in *Trpc4*::Cre⁺ cystitis mice after CNO injection. NS, not significant.

**Extended Data Fig. 8.**
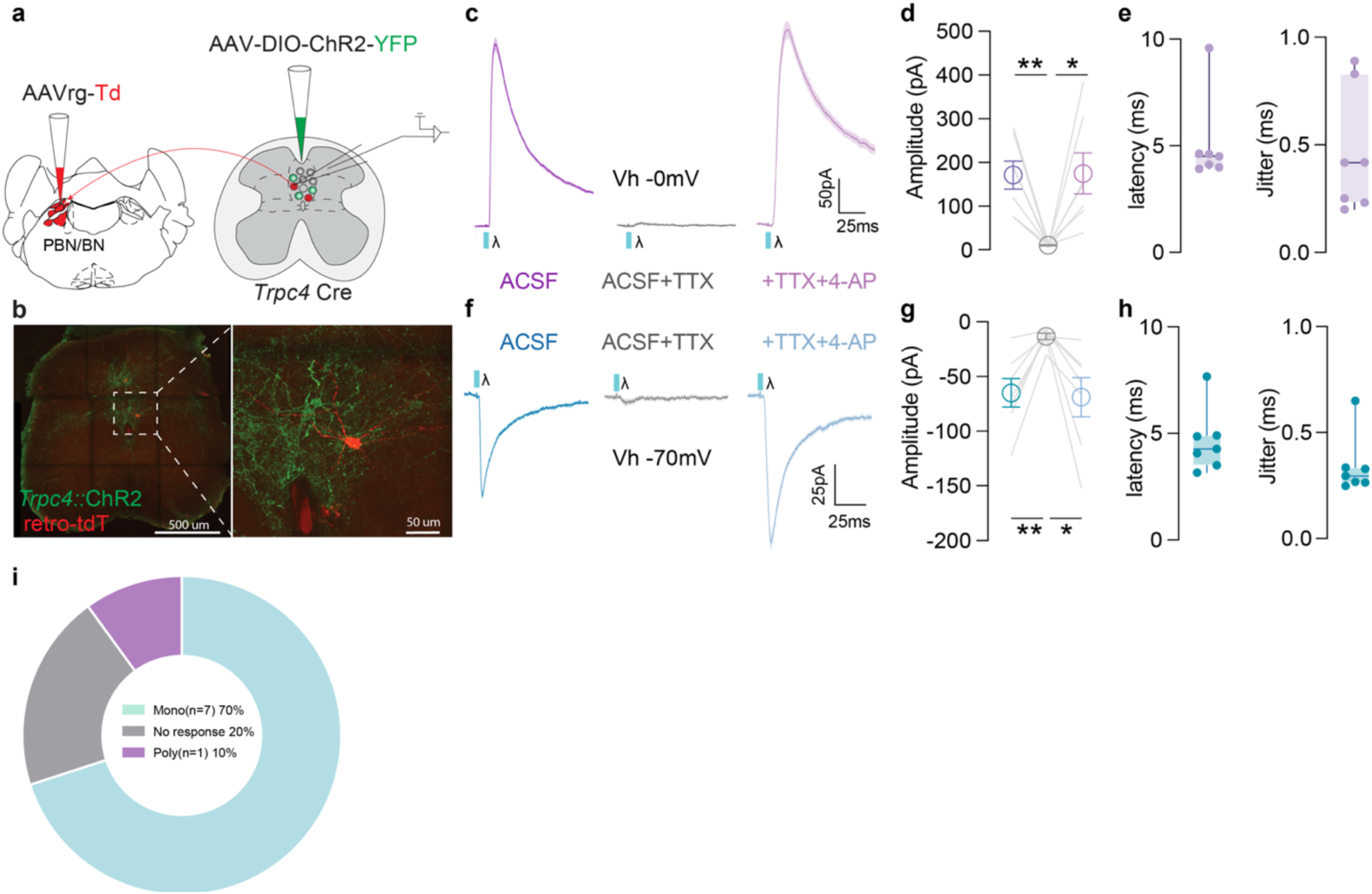
DGC^Trpc4^ interneurons form monosynaptic connections with spinal projection neurons. (a) Experimental strategy combining AAVrg-TdTomato injection into the PBN/BN with AAV-DIO-ChR2-YFP delivery to the spinal cord of *Trpc4*::Cre mice. Scale bar 500 µm (overview), 50 µm (zoom). (b) Representative image showing ChR2-YFP⁺ *Trpc4* neurons and tdTomato-labeled PBN-projecting neurons in the dorsal gray commissure (DGC).(c–e) Light-evoked IPSCs recorded at 0 mV persist in TTX+4AP, confirming monosynaptic inhibitory; quantification of amplitude (d), latency, and jitter (e).(f–h) Light-evoked EPSCs recorded at –70 mV similarly demonstrate monosynaptic excitation input; corresponding quantification of amplitude (g), latency, and jitter (h). (i) Pie chart summarizes postsynaptic responses: 70% monosynaptic, 20% nonresponsive, 10% polysynaptic.

**Extended Data Fig. 9.**
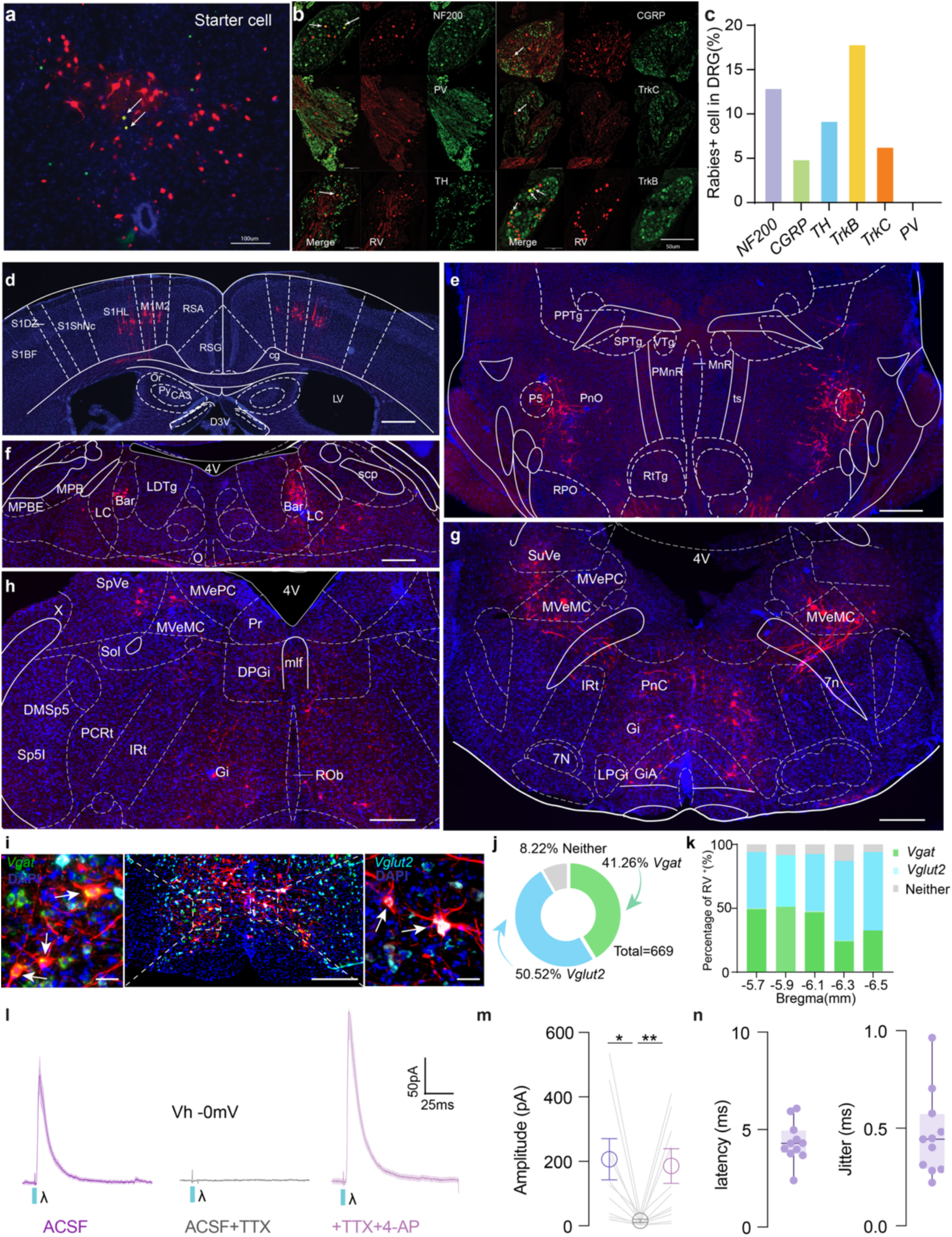
Monosynaptic rabies tracing reveals inputs to DGC*^Trpc4^* neurons. (a) Representative spinal section showing starter cells (TVA-GFP⁺ and N2c G⁺; arrow) in the DGC, scale bar, 100 µm. (b) RV⁺ neurons in lumbosacral DRGs co-express *NF200, CGRP, TH, TrkB, TrkC*, and parvalbumin (*PV*), indicating input from both nociceptive and mechanosensory populations, scale bar, 50 µm. (c) Quantification of molecular marker co-expression among DRG rabies⁺ cells. (d-g) Representative brain sections showing RV⁺ neurons (red) in supraspinal regions, including the motor cortex, pontine nuclei and midbrain, Barrington’s nucleus (Bar), caudal pontine and medullary regions including the gigantocellular nucleus (Gi), pontine reticular nucleus, oral part(PnO) , magnocellular medial vestibular nucleus (MVeMC), parvocellular medial vestibular nucleus (MVePC), spinal vestibular nucleus (SpVe), and superior vestibular nucleus (SuVe). (i) Co-staining *Vgat, Vglut2* demonstrates that supraspinal VMM inputs include both excitatory (*Vglut2*⁺, aqua) and inhibitory (*Vgat*⁺, green) neurons; arrows indicate representative colocalization. (j) Quantification of RV⁺ cells across all levels reveals that 50.52% are *Vglut2⁺*, 41.26% are *Vgat⁺*, and 8.22% are negative for both. (k) Regional distribution of RV⁺ cell phenotypes along the rostrocaudal axis (Bregma −5.7 to −6.5 mm) shows consistent proportions of glutamatergic and GABAergic input across levels. Scale bars: 500 µm (a–e), 50 µm (f left and right), 500 µm (f middle). (l) Light-evoked IPSCs recorded at 0 mV persist in TTX and are recovered with 4AP. (m–n) Quantification of response amplitude, latency, and jitter confirm monosynaptic connectivity.

**Extended Data Fig. 10.**
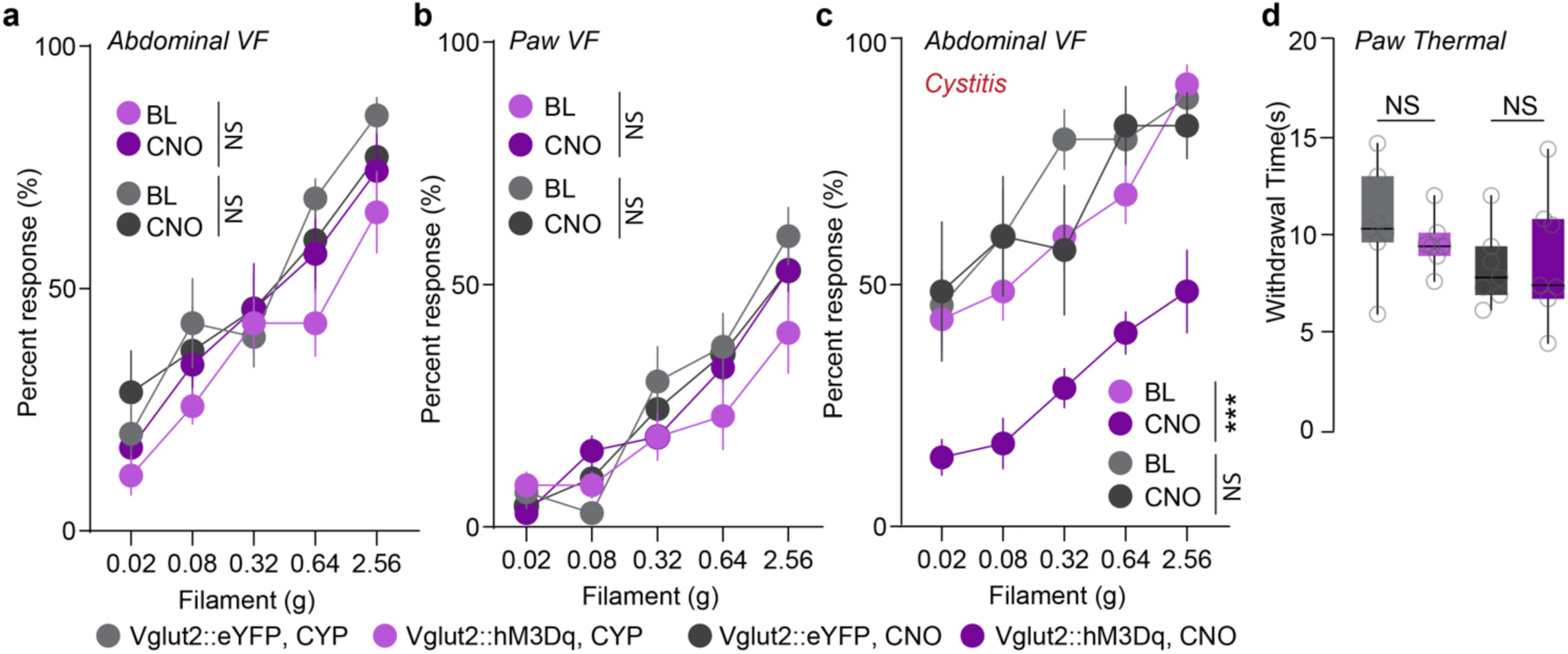
Vglut2⁺ VMM➔DGC pathway selectively attenuates referred sensitivity in cystitis. (a-b) Percent withdrawal responses to abdominal (a) and paw (b) von Frey stimulation show no difference before (BL) and after CNO injection in control (*Vglut2*::eYFP, CYP) and hM3Dq (*Vglut2*::hM3Dq, CYP) groups under normal conditions (P=0.6373, P=0.0948, two-way ANOVA). (c) In cystitis, *Vglut2*::hM3Dq mice show significantly reduced abdominal von Frey responses after CNO treatment compared to BL (right panel, ***P < 0.001, two-way ANOVA). (d)Paw thermal withdrawal latencies are unchanged between groups (P=0.3866, P=0.7448, unpaired t test). NS, not significant.

